# Loss of Arginase 2 Promotes Lung Metastasis in immune-competent hosts via Nitric Oxide Synthase 2-Dependent Th17 Response

**DOI:** 10.64898/2026.06.04.730112

**Authors:** Shu-Ting Chou, Xiaoyong Wang, Jing Yang, Yoonha Hwang, Jiacheng Wang, Yujie Ding, Jeffrey C. Rathmell, Deanna N. Edwards, Jin Chen

## Abstract

Distant metastasis is the leading cause of mortality in many cancers. Although metabolic reprogramming is recognized as a hallmark of cancer, how tumor-intrinsic metabolic enzymes regulate tumor-immune crosstalk during metastatic progression remains poorly understood. Here, using a high-throughput functional CRISPR-Cas9 screen targeting metabolic genes in an orthotopic 4T1 murine mammary carcinoma model of spontaneous lung metastasis, we identify a selective enrichment of arginase 2 (ARG2)-deficient tumor cells in metastatic lungs of immunocompetent but not RAG1-deficient mice, indicating a lymphocyte-dependent mechanism. Loss of ARG2 enhances spontaneous lung metastasis without affecting primary tumor growth. Further, metastatic outgrowth in the lung is not affected when tumor cells are injected intravenously, indicating that ARG2 regulates an early stage of the metastatic cascade. Mechanistically, ARG2 deficiency upregulates nitric oxide synthase 2 (NOS2), resulting in increased nitric oxide production, accumulation of cytosolic DNA, and activation of the cGAS-STING-NF-κB pathway, leading to upregulation of inflammatory cytokines. ARG2-deficient tumors exhibit an immunosuppressive tumor microenvironment characterized by enrichment of Th17 cells and reduced anti-tumor immune populations. Functionally, Th17 cells enhance tumor cell migration *in vitro* and promote spontaneous lung metastasis *in vivo*. Genetic deletion of NOS2 attenuates cytosolic DNA accumulation, reduces STING-NF-κB activation, restores anti-tumor immunity, and suppresses ARG2 deficiency-driven metastatic burden *in vivo*. Collectively, these findings define a tumor cell-intrinsic ARG2-NOS2 axis that regulates inflammatory signaling and the tumor microenvironment to promote metastasis, highlighting a targetable vulnerability in metastatic breast cancer.

## Introduction

Distant metastasis remains the primary cause of mortality in breast cancer, with bone, lung, liver, and brain representing the most frequent metastatic sites across all breast cancer subtypes (1). Although only approximately 6% of patients are diagnosed with distant disease, five-year relative survival rate is markedly reduced compared to localized and regional disease (32.6% vs. 99% and 87.2%, respectively) (2). Metabolic reprogramming is a hallmark of cancer and plays a critical role at multiple stages of metastasis, enabling tumor cells to escape the primary site, survive in circulation, and successfully colonize secondary organs (3–5). While prior studies have established that tumor metabolic alterations confer survival advantages that promote metastatic progression, how tumor-intrinsic metabolic enzymes modulate the interplay between tumor-immune interactions during metastasis remains poorly understood.

Arginine metabolism is a critical regulator of tumor cell fitness and anti-tumor immunity through modulation of polyamine synthesis, nitric oxide (NO) production, and mTOR signaling. L-arginine serves as a common substrate for nitric oxide synthase (NOS), which converts L-arginine into L-citrulline and NO, and for arginase (ARG), which hydrolyzes L-arginine into urea and L-ornithine (6–9). L-ornithine subsequently serves as a precursor for polyamine and proline synthesis or functions as an intermediate in the urea cycle, thereby regulating cell growth and metabolic homeostasis (10). Two arginase isoforms, ARG1 and ARG2, have been identified in mammals. ARG1 is primarily expressed in the cytosol of hepatocytes and myeloid cells, while ARG2 is a mitochondrial enzyme enriched in the kidneys, prostate, T cells, and multiple cancer types (6,7). In cancer, ARG1 has been extensively studied in myeloid and cancer cells as a tumor-promoting factor, while the tumor-intrinsic functions of ARG2 remain less well defined (7). Prior studies have shown that ARG2 promotes tumor proliferation, migration, and immunosuppression in neuroblastoma, melanoma, lung, thyroid, and colorectal cancers (11–15). Conversely, ARG2 suppresses tumor progression in kidney and prostate cancers by inducing polyamine toxicity and depleting the biosynthetic cofactor pyridoxal-5′-phosphate (16,17). Collectively, these findings suggest the pro- or anti-tumor functions of ARG2 may be context dependent and shaped by tumor-specific metabolic programs. However, whether tumor cell ARG2 regulates tumor-immune interactions during metastatic progression remains unknown.

Nitric oxide synthase 2 (NOS2 or iNOS) has emerged as a prognostic marker for cancer, where it modulates tumor growth, inflammatory signaling, and TME remodeling (18,19). Elevated NO levels have been linked to oxidative DNA damage and the release of mitochondrial or nuclear DNA into the cytosol, leading to activation of the cytosolic DNA sensing cGAS-STING pathway (20–26). Downstream activation of NF-κB induces the expression of various inflammatory cytokines, including *Ccl20* and *Il-6* that are key mediators of Th17 cell recruitment and differentiation (27–29). Th17 cells are defined by the lineage-specific transcription factor RORγt and characterized by secretion of cytokines IL-17A, IL-17F, and IL-22 (30). The role of Th17 cells in tumor progression has been paradoxical and appears to depend on tumor type, disease stage, and local microenvironmental cues. In certain contexts, Th17 cells enhance anti-tumor immunity through recruitment and activation of cytotoxic T cells, NK cells, dendritic cells, and neutrophils, whereas in other settings they promote chronic inflammation, immune suppression, epithelial-to-mesenchymal transition (EMT), and metastatic dissemination through IL-17-mediated remodeling of the TME (30–32).

To identify metabolic regulators of breast cancer metastasis that function through modulation of tumor-immune crosstalk, we performed an *in vivo* CRISPR-Cas9 screen targeting metabolic genes in an orthotopic 4T1 murine mammary carcinoma model of spontaneous lung metastasis and identified ARG2 as a metastatic suppressor in immune-competent mice. We show that ARG2 deficiency in breast cancer cells promotes NOS2-dependent chronic inflammatory signaling, leading to activation of the STING-NF-κB axis, increased Th17 infiltration, and establishment of an immunosuppressive TME. These findings demonstrate that ARG2 acts a suppressor of breast cancer metastasis, with its loss driving NOS-mediated inflammatory signaling and Th17 accumulation, offering therapeutic potential for treating metastatic breast cancer.

## Results

### Metabolic CRISPR Screen Identifies Loss of ARG2 as a Driver of Lymphocyte-Dependent Lung Metastasis

To identify key cancer cell-intrinsic metabolic enzymes and nutrient transporters that regulate lung metastasis through modulation of lymphocyte responses, we used an orthotopic 4T1 spontaneous lung metastasis model to perform an *in vivo* metabolic CRISPR screen in immunocompetent and RAG1-deficient mice, which lack mature T and B lymphocytes (Fig. 1A). Cas9-expressing 4T1 cells were transduced with a lentiviral sgRNA library targeting 101 metabolic genes spanning major metabolic pathways (Fig. S1A). To better model clinical management of breast cancer, we orthotopically implanted tumor cells into the mammary fat pads of mice. Primary tumors were surgically resected, and late-stage metastatic lungs were collected for sgRNA quantification by next-generation sequencing (Fig. 1A). Consistent with prior studies demonstrating that *Tsc2* loss promotes metastasis through tumor-intrinsic mechanisms, sgRNAs targeting *Tsc2* were enriched in metastatic lungs from both mouse models (Fig. 1B) (33). Notably, *Arg2*-specific sgRNAs were selectively enriched in metastatic lungs of immunocompetent mice, but not in RAG1-deficient mice or in primary tumors (Fig. 1B-C). These findings suggest that loss of ARG2 promotes lung metastasis in a lymphocyte-dependent manner.

**Figure 1.**
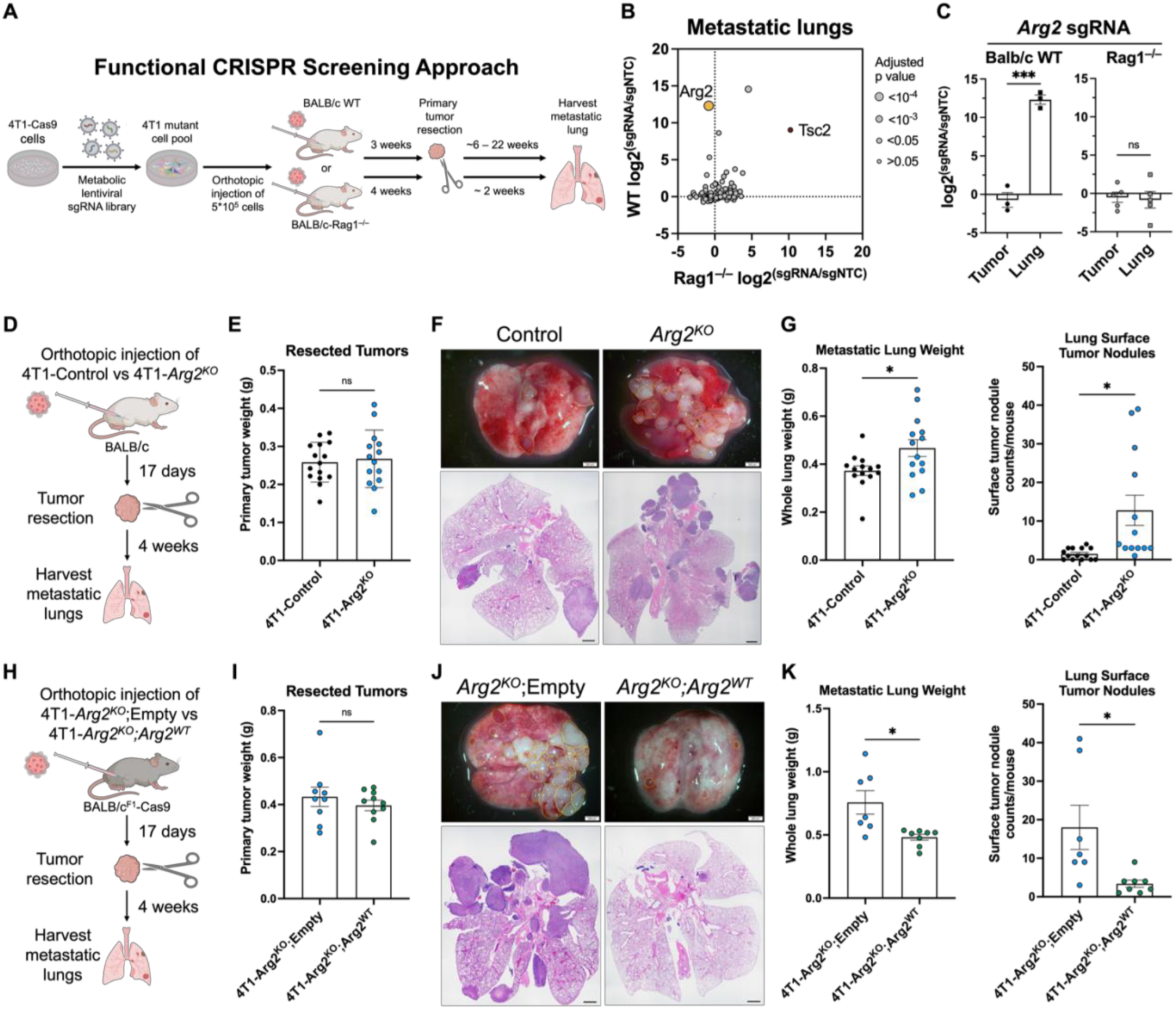
Metabolic CRISPR screen identifies loss of ARG2 as a driver of lymphocyte-dependent lung metastasis. **A-C,** A pooled lentiviral library of 101 sgRNAs targeting metabolic genes was transduced into 4T1-Cas9 cells. Library-expressing cells were orthotopically injected into the mammary fat pad of BALB/cJ wild-type (WT) or *Rag1^−/−^* mice. Primary tumors were resected, allowing for spontaneous metastasis to lung. Metastatic lungs were harvested for next-generation sequencing to identify enrichment or depletion of sgRNAs. **A,** Schematic of the *in vivo* CRISPR screen. **B and C,** Average log_2_ fold change of gene-targeting sgRNAs relative to non-targeting control sgRNAs (sgNTC) in metastatic lungs and primary tumors of BALB/cJ WT (n=3) and *Rag1^−/−^* (n=5) mice. **D-H,** Validation of ARG2 deficiency as a driver of metastasis. 4T1-Control or 4T1-*Arg2^KO^* cells (**D-G**) or 4T1-*Arg2^KO^* cells transduced with a control (4T1-*Arg2^KO^*;Empty) or ARG2 expression vector (4T1-*Arg2^KO^*;Arg2^WT^) (**H-K**) were implanted into the mammary fat pad to allow spontaneous metastasis. **D and H,** Experimental schematics for spontaneous lung metastasis models. **E and I,** Weights of resected primary tumors derived from 4T1 cells (n = 9 to 15 per group). **F and J,** Representative gross lung images and H&E-stained lung sections showing metastatic burden. **G and K,** Whole lung weight and metastatic burden by surface tumor nodule counts of harvested lungs. Data are presented as mean ± SEM and analyzed using Welch’s t-test. *, *p* < 0.05; **, *p* < 0.01; ***, *p* < 0.001. Scale bar is 200 μm for gross lung images and 1 mm H&E-stained lung sections.

To functionally validate the role of ARG2 in lung metastasis, *Arg2* was deleted in 4T1 and EMT6 murine mammary tumor cells using a lentiviral CRISPR-Cas9 system (Fig. S1B). Consistent with prior reports identifying ARG2 as the predominant arginase isoform in breast cancer cells (34), no detectable ARG1 protein expression was detected in either cell line (Fig. S1C). We next evaluated the impact of ARG2 loss on spontaneous metastasis. Control (non-targeting sgRNA) or *Arg2*-knockout (sgArg2, referred to as *Arg2^KO^*) 4T1 cells were orthotopically implanted into the mammary fat pad, followed by primary tumor resection approximately two weeks later (Fig. 1D; Fig. S1D). Loss of ARG2 did not affect primary tumor growth (Fig. 1E), but significantly increased lung metastasis in immunocompetent mice, as evidenced by increased whole lung weight and surface metastatic nodules (Fig. 1F-G). In contrast, enhanced metastasis was not observed in RAG1-deficient mice (Fig. S1E-G), further supporting a lymphocyte-dependent mechanism.

To rule out off-target effects, ARG2 was re-expressed in 4T1-*Arg2^KO^* cells. Western blot analysis confirmed the restoration of ARG2 expression to levels comparable to control cells (Fig. S1H). Re-expression of ARG2 did not alter primary tumor growth but significantly suppressed lung metastasis (Fig. 1H-K), confirming that ARG2 suppresses metastatic progression. To determine whether this phenotype extends across breast cancer models, we utilized the EO771.LMB murine mammary carcinoma line, an EO771 variant that is metastatic to the lung. As EO771.LMB exhibits undetectable endogenous ARG2 expression (Fig. S1I) (35), we engineered EO771.LMB cells to express either wild-type ARG2 (ARG2^WT^) or an enzymatically impaired mutant (ARG2^H160F^) (Fig. S1I-K) (36). Consistently, re-expression of either protein did not alter primary tumor growth (Fig. S1L-M), but expression of ARG2^H160F^ significantly increased lung metastasis (Fig. S1N-O). The differential effects of ARG2^WT^ and ARG2^H160F^ suggest that ARG2-mediated suppression of metastasis is dependent on its ureohydrolase activity. To determine which steps of metastasis are regulated by ARG2, we employed a tail vein injection lung metastasis mode that bypasses early stages of the metastatic cascade (Fig. S1P) (37). In this model, loss of ARG2 did not significantly affect metastatic burden (Fig. S1P-R), indicating that ARG2 primarily regulates early steps of metastasis rather than later colonization or outgrowth in the lung. Together, these findings support a model in which ARG2 enzymatic activity restrains early stages of metastatic progression.

### Loss of ARG2 drives inflammatory gene expression and NOS2 activation

To elucidate the mechanism by which ARG2 loss promotes breast cancer lung metastasis, mRNA sequencing was performed on 4T1-Control and 4T1-*Arg2^KO^*cells. ARG2-deficient cells exhibited upregulation of the nitric oxide synthase gene *Nos2* as well as several inflammatory cytokines and chemokines, including *Ccl20* and *Il-6* (Fig. 2A-B). Indeed, gene set enrichment analysis (GSEA) revealed significant enrichment of inflammatory transcriptional programs in ARG2-deficient cells, including interferon (IFN) and NF-κB signaling pathways (Fig. 2C; Fig. S2A). Notably, elevated expression of *Nos2* and other NF-κB targeted genes, including *Ccl20*, *Cxcl5*, and *Cxcl10* (38–40), was confirmed by qRT-PCR in *Arg2*-deficient cells (Fig. 2D; Fig. S2B), supporting a potential role of these pathways in ARG2-deficient cells.

**Figure 2.**
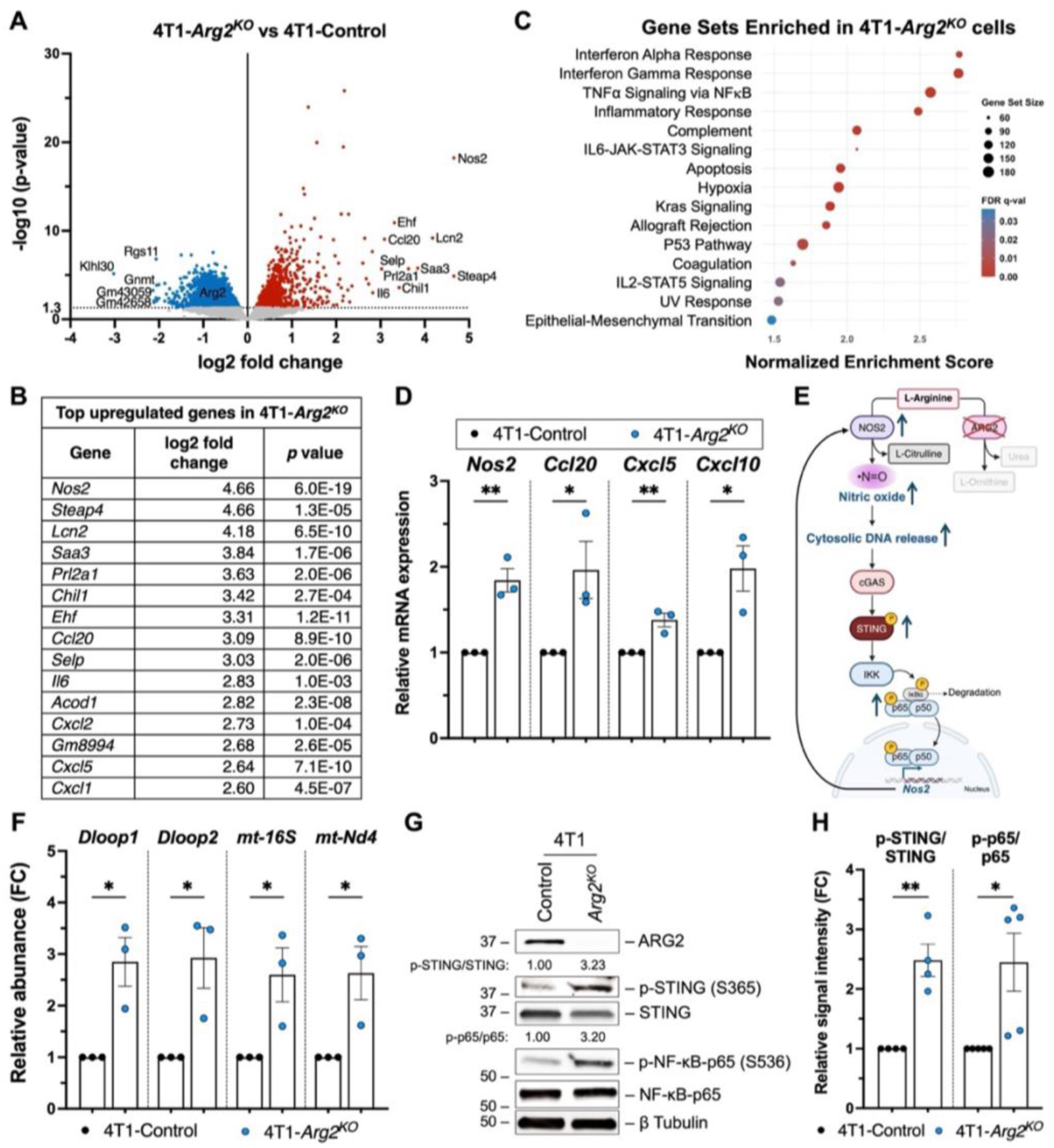
Loss of ARG2 drives NOS2-dependent chronic inflammatory signaling via the STING-NF-κB axis. **A-C,** RNAseq analysis of 4T1-control or 4T1-*Arg2^KO^* cells. **A,** Volcano plot showing log_2_(fold change) versus −log_10_(*p*-value) of genes in 4T1-*Arg2^KO^*cells relative to 4T1-Control cells. Significantly upregulated (red) and downregulated (blue) genes are shown. Gray dots indicate no significant change. **B,** List of the top upregulated genes in 4T1-*Arg2^KO^* cells. **C,** Gene set enrichment analysis (GSEA) showing the top gene sets (Hallmark) enriched in 4T1-*Arg2^KO^* cells compared with control. Gene sets are ranked by normalized enrichment score (NES). Gene set size and false discovery rate (FDR) *q*-value are indicated. **D,** qRT-PCR analysis of NF-κB target genes (*Nos2, Ccl20, Cxcl5,* and *Cxcl10*) in ARG2-deficient (4T1-*Arg2^KO^*) and control (4T1-Control) cells (n=3 per group). **E,** Schematic model illustrating the proposed mechanism by which ARG2 loss activates the NOS2-STING-NF-κB axis. **F,** Mitochondrial DNA (mtDNA) in the cytosolic fraction of 4T1-control or 4T1-*Arg2^KO^* cells (n=3 per group) was determined by qRT-PCR. **G,** Representative western blot analysis of STING and NF-κB p65 activation, including relative densitometry values. **H,** Ratio of phosphorylated protein to total protein was determined for STING and p65 from 4-5 independent blots. Data are presented as mean ± SEM and analyzed using Student’s t-test. *, *p* < 0.05; **, *p* < 0.01.

Metabolomic analysis indicated reduced levels of ornithine, the downstream metabolic product of ARG2 catalytic activity, in *Arg2*-deficient cells (Fig. S2C). However, the levels of the downstream polyamines putrescine, spermidine, and spermine were not significantly changed (Fig. S2C) (41). Additionally, intracellular arginine levels were unchanged, suggesting that L-arginine may be utilized by other enzymes in the absence of ARG2 (Fig. S2C). The nitric oxide synthase NOS2 consumes L-arginine to generate nitric oxide (NO). Given the strong upregulation of *Nos2* in *Arg2*-deficient cells (Fig. 2A-B), we hypothesized that loss of ARG2 increasingly feeds L-arginine to NOS2 thereby enhancing NO production (Fig. 2E). To test this, intracellular NO level was measured using the cell-permeable fluorescent probe 4-Amino-5-Methylamino-2’,7’-Difluorofluorescein Diacetate (DAF-FM DA). 4T1-*Arg2^KO^* cells exhibited significantly elevated NO levels (Fig. S2D). Supplementation with exogenous L-arginine further enhanced NO production in a dose-dependent manner, supporting the notion that L-arginine can be channeled to further fuel NOS2-drive NO production (Fig. S2E).

Elevated NO has been reported to promote oxidative DNA damage, generating cytosolic DNA fragments that drive inflammation (25,26). Therefore, we next assessed whether increased NO production in ARG2-deficient cells was associated with cytosolic DNA accumulation. Subcellular fractionation followed by quantitative PCR revealed significantly increased levels of mitochondrial DNA (mtDNA), including *D-loop1, D-loop2, 16S rRNA,* and *mt-Nd4*, in the cytosolic fraction of *Arg2*-deficient cells, indicating elevated mitochondrial DNA release (Fig. 2F; Fig.S2F). Since cytosolic DNA accumulation activates cGAS-STING signaling (20–23), we next examined activation of the STING-NF-κB pathway. Increased phosphorylation of STING (S365) and NF-κB p65 (S536) was observed in 4T1-*Arg2^KO^* cells relative to controls (Fig. 2G-H), indicating hyperactivation of the STING-NF-κB axis. Collectively, these findings demonstrate that loss of ARG2 promotes *Nos2* upregulation and NO production, cytosolic mtDNA accumulation and activation of inflammatory STING-NF-κB signaling. Given that NOS2 itself is a transcriptional target of NF-κB, these findings support a model in which ARG2 loss establishes a self-sustaining NOS2-STING-NF-κB feedforward loop, resulting in persistent inflammatory signaling and a chronic inflammatory state (42).

### NOS2 Drives Inflammation and Lung Metastasis of ARG2-Deficient Tumor Cells

We next investigated whether NOS2 is required for the enhanced spontaneous lung metastasis induced by ARG2 loss. NOS2 was deleted in 4T1-*Arg2^KO^*cells, and complete ablation of NOS2 protein was confirmed by Western blotting (Fig. S2G). Consistently, DAF-FM DA staining revealed a marked reduction in intracellular NO level in 4T1-*Arg2^KO^;Nos2^KO^* cells relative to *Arg2^KO^* cells (Fig. 3A). Loss of NOS2 also significantly reduced cytosolic mtDNA, indicating a direct link between NOS2-driven NO production and mtDNA release (Fig. 3B; Fig. S2H). We next investigated whether NOS2 contributes to activation of the cGAS-STING-NF-κB pathway and inflammatory gene expression in ARG2-deficient cells. Deletion of NOS2 decreased phosphorylation of STING and NF-κB p65 and expression of NF-κB target genes (Fig. 3C-D), demonstrating NOS2 as a key driver of inflammatory signaling in ARG2-deficient breast cancer cells. To evaluate the functional role of NOS2 in metastasis, *Arg2^KO^*or *Arg2^KO^;Nos2^KO^* cells were orthotopically implanted, and spontaneous lung metastasis was assessed following primary tumor resection (Fig. 3E). Loss of NOS2 significantly suppressed ARG2 loss-driven lung metastasis, as evidenced by decreased whole-lung weight and fewer surface metastatic nodules in metastatic lungs (Fig. 3F-G). Together, these findings demonstrate that NOS2 is a critical mediator of ARG2 deficiency-driven inflammatory signaling and lung metastasis.

**Figure 3.**
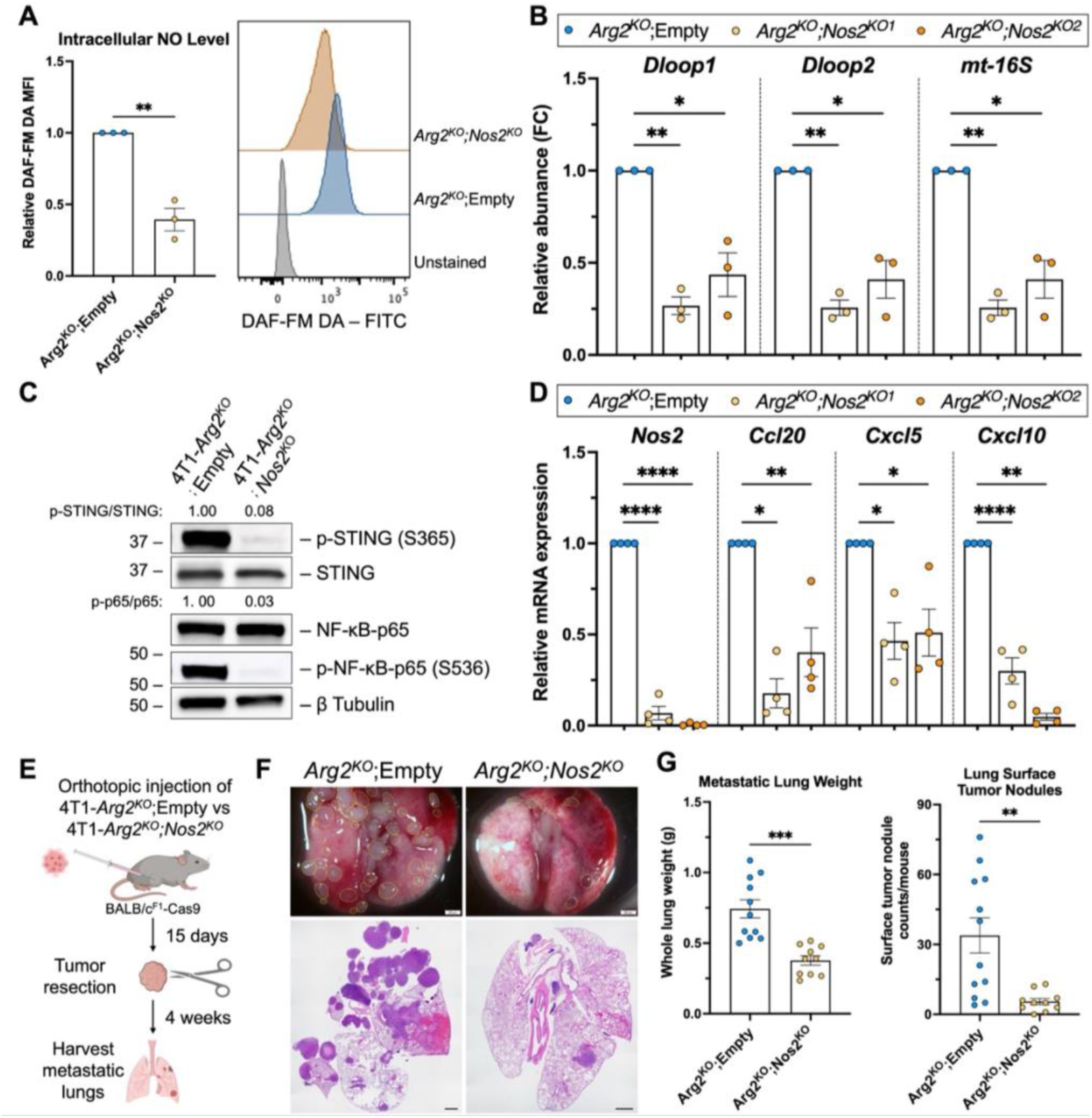
NOS2 drives inflammation and lung metastasis of ARG2-deficient tumor cells. **A,** Flow cytometry analysis of DAF-FM DA fluorescence to assess intracellular NO levels in *Arg2^KO^*;Empty and *Arg2^KO^;Nos2^KO^*4T1 cells. Relative median fluorescence intensity (MFI) is presented (n=3). **B,** mtDNA in the cytosolic fraction of *Arg2^KO^;Empty* and *Arg2^KO^;Nos2^KO^* 4T1 cells was determined by qRT-PCR (n=3). Two independent sgNos2 sequences were evaluated. **C,** Representative western blot analysis of STING and NF-κB p65 phosphorylation in *Arg2^KO^;Empty* and *Arg2^KO^;Nos2^KO^* 4T1 cells, with relative densitometry indicated. **D**, Expression of NF-κB target genes (*Nos2, Ccl20, Cxcl5,* and *Cxcl10*) in 4T1-*Arg2^KO^;Empty* and 4T1-*Arg2^KO^*;*Nos2^KO^* cells (n=4) was determined by qRT-PCR. Two independent sgNos2 sequences were evaluated. **E-G,** The role of NOS2 on metastasis was evaluated by orthotopic implantation of *Arg2^KO^;Empty* and *Arg2^KO^;Nos2^KO^* 4T1 cells in a spontaneous metastasis model. **E,** Experimental schematic is shown. **F**, Representative gross lung images and H&E-stained lung sections showing metastatic burden (n = 10 to 12 per group). **G**, Whole lung weight and surface metastatic tumor nodule counts of harvested lungs. Data are presented as mean ± SEM and analyzed using Welch’s t-test. *, *p* < 0.05; **, *p* < 0.01; ***, *p* < 0.001; ****, *p* < 0.0001. Scale bar is 200 μm for gross lung image and 1 mm H&E-stained lung sections.

### Loss of ARG2 Increases Th17 Cells and Promotes an Immunosuppressive Tumor Microenvironment

Our *in vivo* CRISPR screen and metastasis studies suggested that ARG2 loss enhances lung metastasis through a lymphocyte-dependent mechanism and primarily promotes early metastatic dissemination of the primary tumors by affecting the TME (Fig.1; Fig.S1). We next investigated how ARG2 loss reshapes the immune landscape within primary tumors. Flow cytometry-based immune profiling revealed that ARG2-deficient 4T1 and EMT6 tumors exhibit enrichment of total CD4^+^ T cells and Th17 cells (Fig. 4A; Fig. S3A). The increase in Th17 cells is consistent with the upregulation of *Ccl20* and *Il-6,* which promote Th17 recruitment and differentiation, respectively (Fig. 2A-B, D) (27–29). Additionally, ARG2-deficient tumors displayed an immunosuppressive TME characterized by significant reductions in Th1 cells, effector CD8^+^ IFNγ^+^ T cells, and functional IFNγ^+^ or GZMB^+^ NK cells (Fig. 4B-C; Fig.S3A). These findings suggest that tumor-intrinsic ARG2 expression primarily enriches Th17 cells and creates immune suppression within the primary tumor, potentially impacting early metastatic dissemination.

**Figure 4.**
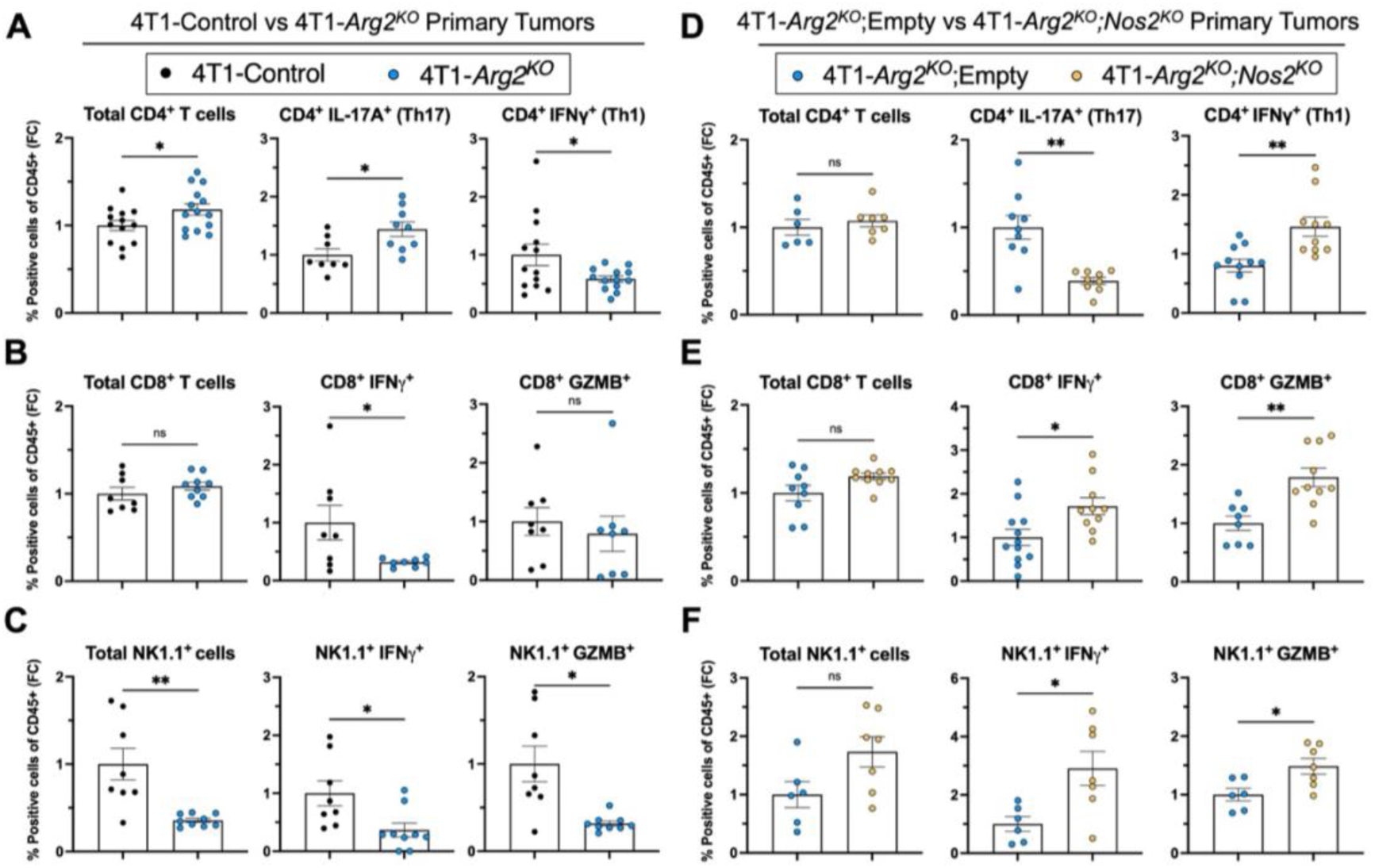
Loss of ARG2 increases Th17 cells and promotes an immunosuppressive tumor microenvironment. 4T1-Control or 4T1-*Arg2^KO^*cells (n=8 to 14 per group) (**A-C**) or 4T1-*Arg2^KO^;*Empty or 4T1-*Arg2^KO^;Nos2^KO^* cells (n=6 to 10 per group) (**D-F**). CD4 T cell populations, including total CD4^+^ cells, CD4^+^ IL-17A^+^ (Th17) cells, and CD4^+^ IFNγ^+^ (Th1) cells (**A and D**), CD8 T cell populations, including total CD8^+^ cells, CD8^+^ IFNγ^+^ cells, and CD8^+^ GZMB^+^ cells (**B and E**), and NK cell populations, including NK1.1^+^ cells, NK1.1^+^ IFNγ^+^ cells, and NK1.1^+^ GZMB^+^ cells (**C and F**) were determined as a percent (%) of CD45^+^ immune cells. Data are presented as fold change (FC) relative to the control. Data are presented as mean ± SEM and analyzed using Welch’s t-test. *, *p* < 0.05; **, *p* < 0.01.

We next assessed whether NOS2 contributes to Th17 enrichment and immune suppression in ARG2-deficient tumors. The *Arg2^KO^;Nos2^KO^* tumors indeed exhibited a significant reduction in Th17 cells (Fig. 4D) compared to the *Arg2^KO^* tumors. Importantly, NOS2 deletion was accompanied by increased Th1 cells and increased effector CD8⁺ T cells and NK cells that are IFNγ^+^ and GZMB^+^ (Fig. 4D-F). Collectively, these findings indicate that loss of ARG2 promotes an immunosuppressive TME through NOS2-mediated inflammatory signaling, characterized by elevated Th17 cells and impaired anti-tumor immune responses.

### Th17 Cells Enhance Tumor Cell Migration and Promote Metastatic Dissemination

To determine whether Th17 cells functionally promote lung metastasis, we co-injected *ex vivo*-differentiated Th17 cells with 4T1 tumor cells into immunocompetent (Fig. 5A-D) and RAG1-deficient mice (Fig. 5E-H). Th17 co-injection did not affect primary tumor growth in either model (Fig. 5B and F). In contrast, Th17 co-injection significantly increased spontaneous lung metastasis in both immunocompetent and RAG1-deficient mice (Fig. 5C-D, G-H), indicating that Th17 cells promote metastatic dissemination. The persistence of enhanced metastasis in RAG1-deficient mice suggests that Th17 cells may promote metastasis, at least in part, through effects on non-lymphocyte populations, including tumor cells.

**Figure 5.**
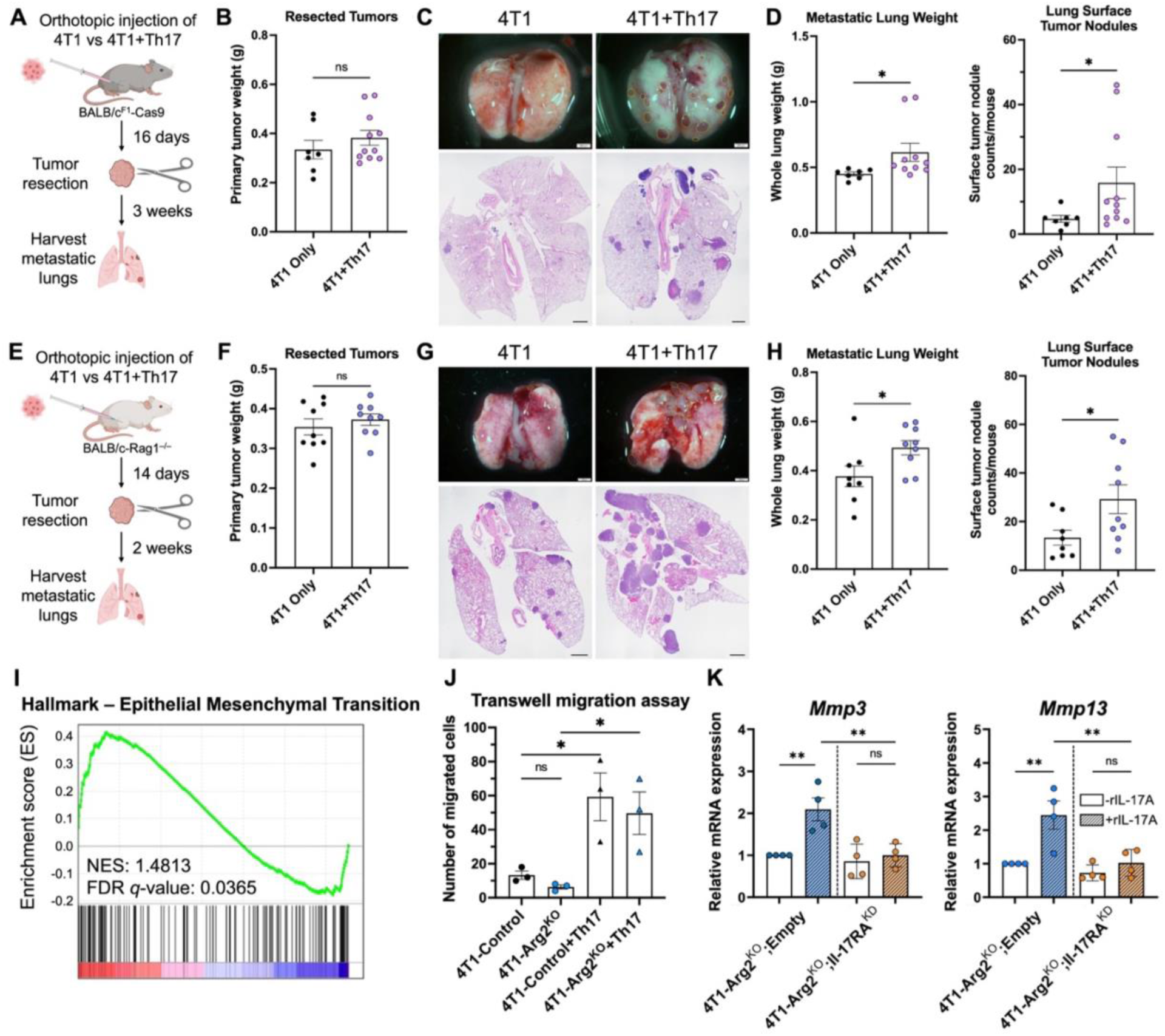
Th17 cells enhance tumor cell migration and promote metastatic dissemination. **A-H,** *In vivo* validation of Th17 cells in promoting spontaneous lung metastasis. 4T1 cells were implanted with (4T1+Th17) or without (4T1 Only) Th17 cells into the mammary fat pad of immunocompetent (**A-D**) and *Rag1^−/−^* mice (**E-H**). **A and E,** Experimental schematics of respective models. **B and F,** Weights of resected primary tumors derived (n = 7 to 11 per group). **C and G,** Representative gross lung images and H&E-stained lung sections showing metastatic burden. **D and H,** Whole lung weight and surface tumor nodule counts of harvested lungs. **I,** Enrichment plot demonstrating epithelial-mesenchymal transition hallmark gene set transcriptome enrichment in 4T1-*Arg2^KO^* cells compared to 4T1-Control cells. Normalized enrichment score (NES) and false discovery rate (FDR) *q*-value is indicated. **J,** Transwell migration assay of 4T1-Control or 4T1-*Arg2^KO^* cells co-cultured with or without *in vitro*-differentiated Th17 cells (n=3 per group). The number of migrated nuclei were quantitated. **K,** qRT-PCR analysis of *Mmp3* and *Mmp13* in 4T1-*Arg2^KO^;Empty* and 4T1-*Arg2^KO^*;*Il-17RA^KD^* cells treated with or without recombinant IL-17A (rIL-17A) (n=4 per group). Data are presented as mean ± SEM and analyzed using Welch’s t-test or ANOVA with Tukey’s post hoc. *, *p* < 0.05; **, *p* < 0.01. Scale bar is 200 μm for gross lung image and 1 mm H&E-stained lung sections.

Due to the lack of a metastatic phenotype in a late-stage model of metastasis of Arg2-deficient tumors and the pro-metastatic role of Th17 cells within the primary tumor (Fig. S1P-R), we next considered the possibility that Th17 cells may induce the migration of breast cancer cells to support metastasis. Indeed, our transcriptome analysis suggest that ARG2-deficient cells exhibit enrichment of EMT-associated genes hallmark, indicating an EMT-like phenotype (Fig. 5I; Fig. S3B). We therefore investigated whether ARG2 loss or Th17 cells directly influence tumor cell migratory capacity using transwell migration assays. While ARG2 deficiency alone did not alter migration, co-culture with Th17 cells significantly increased the number of migrated tumor cells independent of ARG2 status (Fig. 5J; Fig. S3C). These findings suggest that enhanced metastatic dissemination is not driven by intrinsic changes in migratory capacity following ARG2 loss, but rather by Th17-mediated signals.

Since IL-17A is a major effector cytokine produced by Th17 cells that signals through the IL-17RA/RC receptor complex (43), we next examined whether IL-17 receptor signaling regulates tumor cell behavior. Treatment of Arg2-deficient cells with recombinant IL-17A (rIL-17A) induced expression of the metastasis-associated matrix metalloproteinases (MMP), *Mmp3* and *Mmp13*, whereas this induction was attenuated in *Il-17RA*-knockdown (*Il-17RA^KD^*) cells (Fig. 5K; Fig. S3D). These findings are consistent with previous reports linking IL-17 receptor signaling to matrix remodeling and EMT-associated programs (31,32). Collectively, these results demonstrate that Th17 cells directly enhance tumor cell migration and promote metastatic progression, identifying Th17 cells as key mediators of the pro-metastatic phenotype associated with ARG2 deficiency.

### Low ARG2 Expression Predicts Poor Survival and Is Associated with an Increased Th17 Signature in Human Breast Cancer Patients

We have demonstrated that ARG2 loss promotes Th17 cell enrichment and an immunosuppressive tumor microenvironment at the primary tumor site, thereby enhancing tumor cell migration and lung metastasis in syngeneic mouse models. To evaluate the clinical relevance of these findings, we investigated whether ARG2 expression is predictive of outcomes in human breast cancer. Analysis of breast invasive carcinoma datasets revealed that low ARG2 expression was associated with worse overall survival (*p* value = 0.0373, 5-year survival: 62.3% vs 86.0), implicating ARG2 as a determinant of disease progression in advanced breast cancer (Fig. 6A).

**Figure 6.**
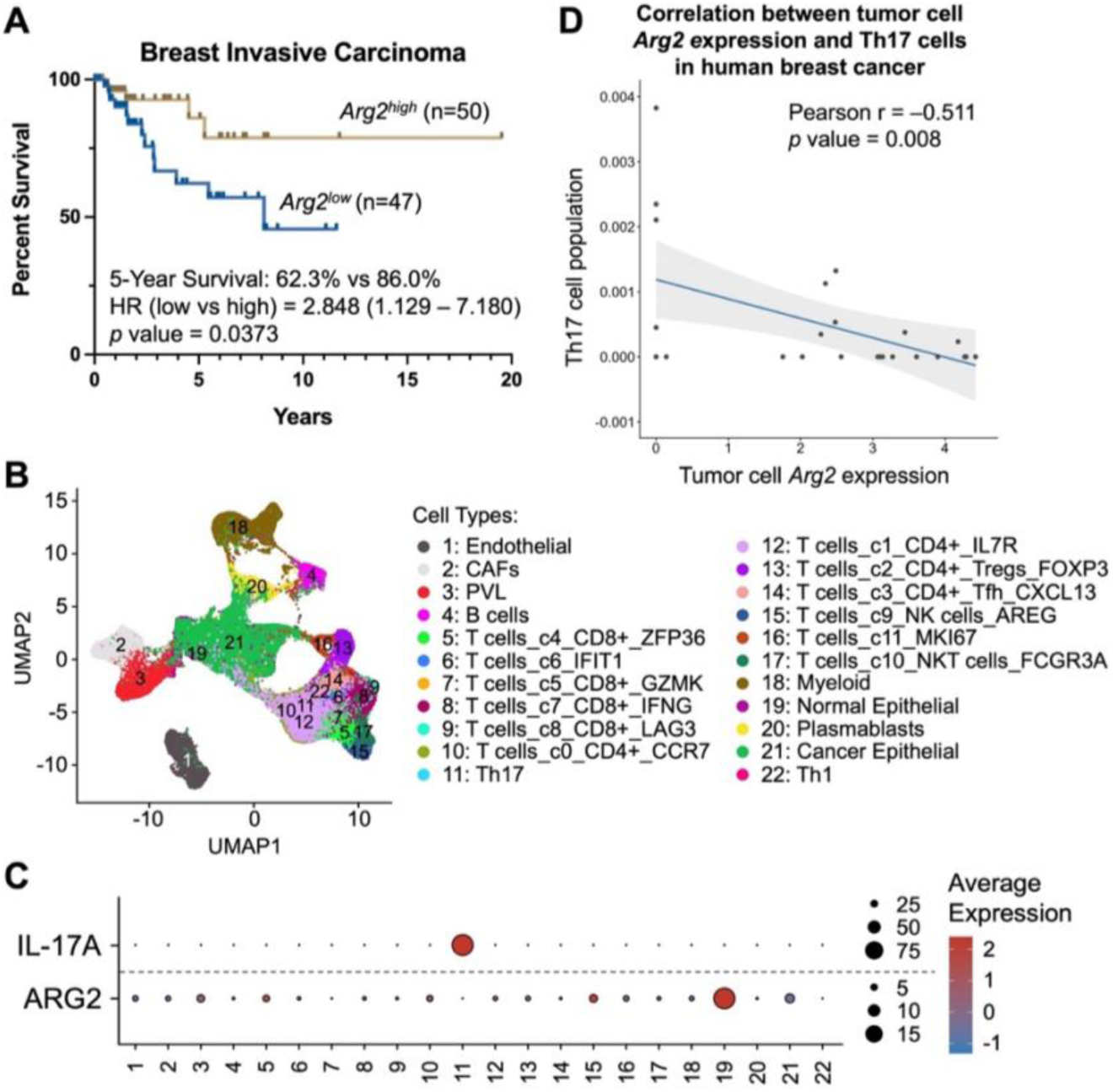
Low ARG2 expression predicts poor survival and is associated with an increased Th17 signature in human breast cancer patients. **A,** Kaplan-Meier analysis of overall survival in patients with breast invasive carcinoma stratified by *Arg2* expression. Five-year survival rates, hazard ratio (HR), and log-rank *p* value are indicated. **B-D,** Analysis of the human breast cancer single-cell RNA sequencing dataset GSE176078. **B,** Uniform Manifold Approximation and Projection (UMAP) visualization of cell populations. **C,** Dot plot showing ARG2 and IL-17A expression across annotated cell populations. Dot size represents the percentage of cells expressing the indicated gene, and color intensity indicates average expression level. **D,** Pearson correlation analysis between ARG2 expression in tumor epithelial cells and the abundance of Th17 cells across human breast cancer samples. Each symbol represent an individual patient. Shaded area represents the 95% confidence interval.

To determine whether the inverse relationship between ARG2 expression and Th17 responses observed in murine models is also present in human breast cancer, we analyzed a single-cell transcriptomic dataset of 26 breast tumors from patients (Fig. 6B) (44). ARG2 expression was primarily detected in normal epithelial cells and T cells, whereas malignant epithelial cells exhibited relatively low ARG2 expression (Fig. 6C). In contrast, IL-17A expression was largely restricted to CD4⁺ T cells, consistent with a Th17 population (Fig. 6C). Correlation analysis revealed a significant inverse correlation between ARG2 expression in tumor cells and Th17 cell abundance (r = -0.511, p = 0.008) (Fig. 6D). Collectively, these findings demonstrate that reduced ARG2 expression is associated with poor clinical outcomes in breast cancer patients and support a potential link between ARG2 loss and enhanced Th17-mediated response in human tumors. These clinical observations are consistent with our mechanistic studies in mouse models and suggest that the ARG2-NOS2-Th17 axis may contribute to metastatic progression in human breast cancer.

## Discussion

Distant metastasis remains the leading cause of mortality in breast cancer, and increasing evidence suggests that, beyond supporting tumor cell growth, metabolic reprogramming promotes metastatic progression by shaping the TME through modulation of immune and stromal cell function, thereby driving tumor invasion, matrix remodeling, and EMT (3–5,45,46). Using an *in vivo* CRISPR-Cas9 screen targeting key metabolic pathways, we identified ARG2 as a suppressor of breast cancer metastasis that functions through regulation of tumor-immune interactions rather than primary tumor growth. Loss of ARG2 selectively enhanced spontaneous lung metastasis in immunocompetent but not lymphocyte-deficient hosts, revealing a lymphocyte-dependent mechanism. Mechanistically, ARG2 deficiency increased NOS2-dependent NO production, leading to activation of the cGAS-STING-NF-κB pathway, upregulation of inflammatory cytokines, and enrichment of Th17 cells. These changes are associated with a remodeled TME that favors an immunosuppressive state, thereby promoting metastatic dissemination. Consistent with these findings, genetic deletion of NOS2 restored anti-tumor immunity and suppressed metastasis, identifying the ARG2-NOS2-Th17 axis as a key regulator of metastatic progression.

ARG2 catalyzes the hydrolysis of L-arginine into urea and L-ornithine, the latter of which can be utilized for proline and polyamine synthesis or function as an intermediate in the urea cycle. Polyamines have been implicated in cytotoxicity against renal cell carcinoma (17), as well as regulation of anti-tumor T cell responses (47–49). However, our metabolomics data revealed that loss of ARG2 in tumor cells decreased ornithine level but polyamines, including putrescine, spermidine, and spermine, are unchanged (Fig. S2C), suggesting that polyamines do not play a predominant role in our breast model. While prior studies have also attributed ARG2-mediated depletion of extracellular L-arginine to immune suppression (7), we found that arginine levels remain the same between control and ARG2-deficient cells (Fig. S2C), suggesting that L-arginine is metabolized via an alternative pathway. Indeed, our data suggest that L-arginine is converted by NOS2 to produce nitric oxide, leading to inflammatory cytokine production that remodels tumor microenvironment, enhancing Th17 infiltration and reducing Th1, effector CD8+ T cells, and functional NK cell responses (Fig. 3 and 4). In particular, our data point toward a Th17-dependent impact toward increased lung metastasis (Fig. 5). These findings identify a previously unrecognized mechanism by which ARG2 shapes anti-tumor immunity.

The effects of Th17 cells in cancer are pleiotropic, as they can promote angiogenesis, tumor growth, metastasis, chronic inflammation, and immune suppression in some contexts while enhancing effector T cell recruitment and anti-tumor immunity in others (30). In our model, increased intratumoral Th17 infiltration was associated with diminished anti-tumor immunity and enhanced metastatic progression. Importantly, ARG2 deficiency alone did not enhance tumor cell migration *in vitro* despite enrichment of a EMT-associated gene signature, indicating that the increased metastatic dissemination observed *in vivo* is not solely attributable to tumor-intrinsic changes. Rather, Th17 cells directly promoted tumor cell migration *in vitro* and increased metastatic burden *in vivo*, even in lymphocyte-deficient hosts, establishing a functional role for Th17 cells in driving metastasis. Given that Th17 cells can secrete IL-22 in addition to IL-17, one possible mechanism for Th17-induced metastasis is through inhibition of NK cells by IL-22, consistent with a previous report (50). Alternatively, remodeling of TME by loss of ARG2 resulted in decreased CD8 T cells effector function (Fig. 4), which could contribute to increased metastasis. However, promotion of lung metastasis in Rag-1 null animals by Th17 co-injection established Th17 cells as the key pro-metastatic downstream effectors of ARG2-NOS2-driven inflammatory signaling.

Our findings are supported by analyses of human breast cancer datasets demonstrating that low ARG2 expression is associated with poor survival in advanced disease. Moreover, we observed an inverse correlation between tumor ARG2 expression and Th17 cell abundance in patient tumors (Fig. 6), suggesting that the mechanisms identified in murine models may be conserved in humans and supporting a clinically relevant link between ARG2 loss and Th17-mediated immune remodeling. The identification of the ARG2-NOS2-Th17 axis as a driver of metastatic progression provides multiple potential therapeutic intervention points. Inhibition of NOS2 or downstream inflammatory signaling, such as cGAS-STING, may restore anti-tumor immunity and suppress metastatic progression. Alternatively, targeting Th17-associated cytokines or inhibition of RORγt transcription factor may mitigate pro-metastatic immune remodeling and reduce tumor invasiveness. Collectively, these findings establish the ARG2–NOS2–Th17 axis as a mechanistic link between tumor metabolism, inflammatory TME remodeling, and metastatic progression, highlighting a potential therapeutic vulnerability in metastatic breast cancer.

## Materials and methods

### Mice

All experiments were performed at Vanderbilt University in accordance with Institutional Animal Care and Utilization Committee–approved protocols and conformed to all relevant regulatory standards. All mice were housed in a pathogen-free animal facility, and immunocompromised RAG1-deficient mice were housed in sterile housing. BALB/cJ (#651), C57BL/6J (#664), C57BL/6-Cas9 (#26179) and BALB/cJ-*Rag1*^−/−^ (#3145) mice were purchased from Jackson Laboratory. For immunological studies using BALB/c-derived genetically modified 4T1 and EMT6 cell lines that express Cas9, Cas9-expressing BALB/c^F1^-Cas9 mice were generated by crossing BALB/cJ and C57BL/6-Cas9 mice. F1 generation progeny were used for experiments, as indicated. Eight to twelve-week-old female mice were used for all animal experiments.

### Cell culture

The murine breast cancer cell lines 4T1, EMT6, and E0771.LMB were purchased from American Type Culture Collection (ATCC). All tumor cell lines were maintained at low passages and cultured in Dulbecco’s modified Eagle’s medium (DMEM) supplemented with 10% fetal bovine serum and 100 U/ml penicillin/streptomycin mixture in a humidified atmosphere at 37°C with 5% CO_2_.

### Constructs and lentiviral transduction

Lentivirus was produced by transfecting HEK293FT cells with indicated lentiviral expression plasmids, pCMV-VSV-G, and psPAX2 using the Lipofectamine 3000 transfection reagent. The virus supernatant was harvested 48 hr after transfection, filtered through a 0.45 μM filter, and polybrene was added at 20 μg/ml final concentration before transduction. Prior to viral transduction, 2.5×10^5^ cells were plated into 6-well plates 24 hr before. For viral transduction, cells were incubated in media containing virus and polybrene for 72 hr. Cells were allowed to recover for 24 hr before antibiotic selection. For antibiotic selection, transduced cells were incubated in puromycin- or blasticidin S HCl containing media at 2 μg/ml or 4 μg/ml, respectively, and cultured for three days.

Lentiviral vectors plentiCRISPRv2-Puro, plentiCRISPRv2-Blast, and plenti-CMV-Blast were obtained from Addgene. For genetic knockout using CRISPR-Cas9, sgRNAs were cloned into plentiCRISPRv2-Puro or plentiCRISPRv2-Blast vectors. For *Arg2* re-expression, gBlocks (Integrated DNA Technologies) double-stranded DNA fragments encoding *Arg2^WT^* or mutant *Arg2^H160F^* were cloned into the mammalian expression vector plenti-CMV-Blast. Indel PCR was performed to verify the sequence and orientation of the inserted sgRNAs and gBlock DNAs. Sequences of the sgRNAs, gBlock DNA fragments, and indel PCR primers are listed in Supplementary Table 1.

### *In vivo* CRISPR Screening

The metabolic sgRNA library was curated by referencing the Mouse CRISPR Knockout Pooled Library (Brie) (Addgene, Pooled Library #73632) targeting metabolic enzymes involved in lipid metabolism, TCA cycle, oxidative phosphorylation, amino acid metabolism, glycolysis, and nucleic acid metabolism. The library contained four sgRNAs targeting each gene and eight non-targeting control sgRNAs. sgRNAs targeting *Tsc2* served as positive control. The sgRNA library was cloned into the lentivirus expression vector plentiCRISPRv2-Puro, and the resultant plasmid pool was transfected into HEK293FT cells to generate the metabolic lentiviral sgRNA library. 4T1 cells expressing Cas9 were transduced with the lentiviral sgRNA library. Five hundred thousand cells from the 4T1 mutant cell pool were injected orthotopically into the mammary fat pad of female BALB/cJ wild-type or RAG1-deficient mice, and primary tumors were resected three and four weeks post-inoculation, respectively. Late-stage metastatic lungs were harvested at endpoint of studies.

Genomic DNA from plasmids and cell injection controls were purified using the PureLink™ Genomic DNA Mini Kit following manufacturer’s instructions. For tumors and metastatic lungs, tissues were minced by scissors and lysed in 12 ml of NK lysis buffer (50 mM Tris-HCl, 50 mM EDTA, 1% SDS, pH 8.0) supplemented with proteinase K (20 mg/ml) at 55°C overnight. After recovering to room temperature, RNaseA (100 mg/ml) was added then incubated at 37°C for 30 min. To precipitate proteins, 4 ml of pre-chilled 7.5M ammonium acetate was added, samples were mixed by vortexing and centrifuged at 4,000 x g for 10 min. Supernatant was collected, 12 ml of 99.99% isopropanol was added, and samples were mixed by inversion followed by centrifugation at 4,000 x g for 10 min. The resulting DNA pellet was washed by 70% ethanol then centrifuged. After air-drying the pellet at room temperature for 30 min, 500 μl of 1X TE buffer was added, samples were incubated at 65°C for 1hr, then place at room temperature for overnight. The sgRNA sequences were amplified from genomic DNA samples by two rounds of PCR: first round with Adaptor primers and second round with barcoded Illumina sequencing primers. The amplicons were then purified by agarose gel extraction, combined at equimolar concentrations, and sequenced for 150 cycles in paired-end mode on the Illumina Novaseq 6000 platform at the Vanderbilt Technologies for Advanced Genomics. At least 1000-fold representation of the library was maintained throughout the process. FASTQ files were analyzed using the Model-based Analysis of Genome-wide CRISPR-Cas9 Knockout (MAGeCK v0.5.0.3) method for statistical analysis.

### Mouse tumor and metastasis studies

For orthotopic mammary fat pad injections, cells were embedded in 100 μl of Matrigel Growth Factor Reduced (GFR) Basement Membrane Matrix (Corning #354230). For 4T1 and E0771.LMB spontaneous lung metastasis experiments, 3×10^5^ or 2.5×10^5^ cells were injected orthotopically into BALB/cJ or BALB^F1^-Cas9 mice. For EMT6 tumor studies, 2.0×10^5^ cells were injected orthotopically into BALB^F1^-Cas9 mice. For 4T1 and Th17 co-injection experiments, 1.5×10^5^ of tumor cells and 1.5×10^5^ of RORγt^+^ CD4^+^ T cells were injected into BALB/cJ-*Rag1*^−/−^ or BALB^F1^-*Cas9* mice. The length (L) and width (W) of the primary tumors were measured using calipers. Tumor size was calculated using the following formula: LxW^2^xπ/6. Primary tumors were resected when tumors in all mice reached 300 to 500 mm^3^ in size on approximately day 14 to 17 post-injection. For tail-vein injections, 2×10^5^ 4T1 cells in 200 μl of PBS were injected into the tail-vein of BALB^F1^-Cas9 mice. Late-stage metastatic lungs were harvested at a single time point, approximately three to four weeks post-resection when mice exhibit signs of illness and were fixed in 10% neutral buffered formalin. H&E staining of metastatic lungs was performed by the Translational Pathology Shared Resource at Vanderbilt University Medical Center.

### RNA sequencing

RNA was purified from 4T1-*Control* and 4T1-*Arg2^KO^*cells using the RNeasy Plus Mini Kit (Qiagen #74104) according to the manufacturer’s instructions. RNA quantity and quality were assessed using the Qubit High-Sensitivity RNA Assay Kit (Thermo Fisher #Q32852). RNA sequencing was performed by Azenta using the Illumina Novaseq platform. Sequencing reads were trimmed and mapped to the mouse reference genome GRCm38 using STAR aligner (v2.7.9a). Gene-level counts were generated using the SummarizeOverlaps function from the GenomicAlignments R package. Differential gene expression analysis between groups was performed using DESeq2 (v1.24.0). Adjusted p values were calculated using the Benjamini–Hochberg method to control the false discovery rate (FDR). Log_2_ fold change (FC) values were computed relative to controls.

### Western blot analysis

Cells were harvested in RIPA buffer (Sigma Aldrich #R0278) containing Roche cOmplete, Mini Protease Inhibitor Cocktail (Sigma Aldrich #04693124001) and Roche PhosSTOP (Sigma Aldrich #4906845001) at approximately 50 μl RIPA buffer per one million cells. Cell lysates were centrifuged at 14,000 x g for 10 min at 4°C to pellet debris and supernatants were collected. Protein lysates were quantified by DC Protein Assay (Bio-Rad #5000112) using bovine serum albumin (BSA) as protein standard. Protein lysates were resolved by Tris-Glycine SDS-PAGE and were transferred to 0.2 μm nitrocellulose membranes at 2.5A, 25V for 10 min using the Trans-Blot Turbo Transfer System. Membranes were blocked using the Intercept TBS (Thermo Fisher #92760001) or PBS Blocking Buffer (Thermo Fisher #92790001) at room temperature for 2 hr and then incubated with the indicated primary antibodies diluted in TBST (20 mM Tris, 135 mM NaCl, and 0.02% Tween 20) supplemented with 1% BSA. Primary antibodies were detected with horseradish peroxidase-linked secondary antibodies followed by exposure to Clarity Western ECL Substrate (Bio-Rad #1708891) or SuperSignal West Femto Maximum Sensitivity Substrate (Thermo Fisher #34095). Relative protein levels were quantified by measuring band intensities using ImageJ software with background subtracted and normalized to the loading control β-tubulin. Antibodies used for western blot analysis are listed in Supplementary Table 2.

### Quantitative reverse transcription PCR

Total RNA was isolated using the RNeasy Plus Mini Kit and cDNA was synthesized using the iScript cDNA Synthesis Kit (Bio-Rad #1708891) with 1000 ng of input total RNA per reaction. qRT-PCR was performed using the iTaq Universal SYBR Green Supermix (Bio-Rad #1725121) with 1:10 diluted cDNA samples and primer pairs at 500 nM per primer. Cycling conditions were 10 min at 95°C, followed by 40 cycles of 15s at 95°C and 1 min at 60°C in the QuantStudio 3 Real-Time PCR System. mRNA expression was normalized to an endogenous housekeeping gene ribosomal protein L37 *(Rpl37)* and relative expression was analyzed using the 2^−ΔΔCT^ method. Gene-specific primers were purchased from Integrated DNA Technologies and sequences are listed in Supplementary Table 1.

### Flow cytometry analysis

For resected primary tumors and metastatic lungs, tissues were dissociated in RPMI-1640 with 5% FBS, collagenase IA (1 mg/mL; Sigma Aldrich #C9891), and DNase I (0.25 mg/mL; Sigma Aldrich #DN25) using the gentleMACS™ Octo Dissociator with Heaters system (Miltenyi Biotec) with the 37C_m_TDK_2 and 37C_m_LDK_1 programs, respectively. To obtain single-cell suspensions, digested tissues were filtered through a 70 μm cell strainer and red blood cells were lysed using ACK lysis buffer (KD Medical #RGF-3015). Cells were stained for viability with Ghost Dye Violet 510 at a 1:1000 dilution in PBS for 15 min at 4°C. Following blocking with the anti-CD16/32 mouse Fc block (Cytek Biosciences #70-0161) for 10 min at room temperature, extracellular staining was performed using fluorophore-conjugated antibody solution in PBS containing 5% FBS and 0.1% sodium azide for 30 min at 4°C. Intracellular cytokine staining of IFNγ, TNFα, GZMB, and IL-17A was performed using the BD Cytofix/Cytoperm Fixation/Permeabilization Kit (BD Biosciences #554714) following manufacturer’s instructions. Stained cells were analyzed on the BD Fortessa using the BD FACSDiva software and data were analyzed with FlowJo. Antibodies used for flow cytometry analysis are listed in Supplementary Table 2.

### DAF-FM DA staining for intracellular NO detection

Cells were detached using 0.25% Trypsin-EDTA and washed three times with PBS. Cells were resuspended in 5 μM DAF-FM DA in XF DMEM Medium, pH 7.4 (Agilent Technologies #10-357-5100) and incubated at 37°C for 20 min. After washing twice with PBS, cells were analyzed immediately by flow cytometry at the FITC channel.

### Cytosolic DNA extraction and quantitative PCR of mitochondrial DNA

Cells were lysed in digitonin buffer containing 20 μg/ml digitonin, 150 mM NaCl, and 50 mM HEPES on an end-over-end tube rotator for 10 min at 4°C. The cytosolic supernatant was collected, and both the cell pellet and a small portion of the cytosolic fraction were used for protein immunoblotting to assess fraction purity. DNA isolation from the cytosolic fractions was performed using the PureLink™ Genomic DNA Mini Kit (Thermo Fisher #K182001) following manufacturer’s instructions. Quantitative PCR of mitochondrial DNA in the cytosol was performed using the iTaq Universal SYBR Green Supermix with 4 ng of DNA input and primer pairs at 500 nM per primer. Primers were purchased from Integrated DNA Technologies and sequences are listed in Supplementary Table 1.

### *In vitro* Th17 differentiation

Murine CD4^+^ T cells were isolated from the spleens and lymph nodes using the Mouse Naive CD4^+^ T Cell Isolation Kit (Miltenyi Biotec #130-104-453) according to the manufacturer’s instructions. Naive CD4^+^ T cells were activated using plate-bound anti-mouse CD3 (2 μg/mL; Cytek Biosciences #70-0032) and anti-mouse CD28 (2 μg/mL; Cytek Biosciences #70-0281) antibodies at 1 million cells/well in 12-well plate. Cells were cultured for 3 days at 37°C with 5% CO_2_ with Th17-skewing cytokines and antibodies: mouse IL-6 (60 ng/ml; Miltenyi Biotec #130-096-685), human TGFβ (1.5 ng/ml; Thermo Fisher #100-21C), mouse IL-23 (20 ng/ml; Miltenyi Biotec #130-096-677), mouse IL-1β (10 ng/ml; Miltenyi Biotec #130-101-681), anti-mouse IL-4 (10 μg/ml; BioXCell #BP0045), and anti-mouse IFNγ (10 μg/ml; BioXCell #BP0055) prepared in RPMI 1640 media supplemented with 10 mM HEPES, 50 μM 2-mercaptoethanol, 100 U/ml penicillin/streptomycin, and 2 mM glutamine. Differentiated Th17 cells were stained with viability dye Ghost Dye Violet 510, stained for CD4 and TCRβ, then fixed with 4% paraformaldehyde. Transcription factor staining of RORγt was performed using the FoxP3/Transcription Factor Staining Buffer Kit (Cytek Biosciences #TNB-0607) following manufacturer’s instructions, and stained cells were analyzed by flow cytometry analysis.

### Arginase activity assay

Arginase activity was determined using the QuantiChrom Arginase Assay Kit (BioAssay Systems #DARG-100) following manufacturer’s protocol. Briefly, cells were lysed in 0.4% Triton X-100 lysis buffer (16 mM HEPES, pH 7.4; 0.8 mM EDTA; 4 mM pyrophosphate; 4 mM glycerophosphate; 0.4% Triton X-100 in 1X PBS containing protease and phosphatase inhibitors on ice for 10 min. Cell lysates were centrifuged at 14,000 x g for 10 min at 4°C and supernatants were collected. Lysate supernatants were incubated with Arginase Reaction Buffer at 37°C for 2 hr. An urea standard curve was made, and samples were mixed with Urea Reagent buffer and incubated for 1 hr at room temperature. The absorbance was read at 430 nm to quantify urea production and data were normalized to protein concentration in each sample determined using the DC Protein Assay, as described above.

### Transwell migration assay

The effect of Th17 on tumor cell migratory capacity was measured on 6.5 mm diameter transwell filters with an 8 μM pore size (Corning #3422). Trypsinized tumor cells (2.5×10^4^ cells) with or without in vitro differentiated Th17 (2.5×10^4^ cells) were washed with PBS, resuspended in 100 μl of RPMI-1640 containing 10% FBS, and placed into the Transwell insert. The lower chamber was filled with 600 μl of RPMI-1640 containing 10% FBS. After 24 hr of incubation at 37°C with 5% CO2, cells on the upper chamber were removed with a cotton swab and filters were washed twice with PBS. Cells present on the lower surface of the filters were fixed in 4% paraformaldehyde for 10 min, then stained with NucBlue Fixed Cell ReadyProbes Reagent (DAPI; Thermo Fisher #R37606) for 10 min. Filters were then washed twice with PBS and cut from the inserts. The nuclei from five different randomly selected fields of each Transwell were imaged at 10X magnification using a fluorescence microscope and total nuclei were counted as a measure of migrated cells. The experiment was repeated three times, and results were averaged.

### Metabolomics analysis

Metabolomic profiling was performed by Metabolon on cell pellets collected from control (sgNTC) and *Arg2^KO^* 4T1 cells (n=6 per group) using ultra-high-performance liquid chromatography-tandem mass spectrometry (UPLC-MS/MS), as previously described (51). Metabolites were identified through comparison with authenticated standards based on retention index, mass spectral matching, and score matching. A total of 837 metabolites were detected and normalized to protein content. Normalized metabolite data were subsequently organized into major metabolic classes and subclasses for downstream analyses.

### Single-cell RNA sequencing analysis

Publicly available single-cell RNA sequencing data from human breast cancer samples (GSE176078) (44) was downloaded from the Gene Expression Omnibus (GEO) and analyzed using Seurat in R. Cell-type annotations provided by the original study were used to identify malignant tumor cells and immune cell populations. ARG2 expression was evaluated across annotated cell types and visualized using UMAP and dot plots. For correlation analysis, the expression of ARG2 in tumor cells and frequency CD4^+^ IL17A^+^ cells were calculated for each patient sample, and associations were assessed using Pearson correlation analysis.

### Statistical analysis

Statistical analyses were performed using GraphPad Prism or R Studio. Data are presented are demonstrated as means ± SEMs. Each symbol represents an individual mouse or independent biological replicate. For two-group analyses, Student’s t-test or Welch’s t-test were performed to compare sample means. For multiple-group analyses, one-way analysis of variance (ANOVA) accompanied by a post hoc Tukey test was used. Outliers were excluded based on the ROUT method with Q value set at 1%. All *p* values are two-tailed and *p* < 0.05 was considered to represent statistically significant events. Significance was recorded as *, *p* < 0.05; **, *p* < 0.01; ***, *p* < 0.001; ****, *p* < 0.0001. Kaplan-Meier survival analysis of overall survival was performed using the breast invasive carcinoma datasets from The Cancer Genome Atlas (TCGA) available through the Human Protein Atlas (www.proteinatlas.org) (52). Patients were stratified into ARG2-high and ARG2-low groups based on the top and bottom 20% of ARG2 expression levels, respectively.

## Acknowledgements

The authors would like to thank Holly Algood (Vanderbilt University Medical Center) for thoughtful discussions on Th17 cells. We also thank Wentao Deng (Vanderbilt University Medical Center) and Channing Chi (Vanderbilt University) for their assistance performing and analyzing the CRISPR screen. This work was supported by NIH grants CA250506 and CA271176, a VA Career Scientist Award (5IK6BX005391), and a VA Merit Award (5101BX000134) to JC. DNE is supported by a Department of Defense CDMRP grant W81XWH2210109 and a Career Enhancement Award, funded through the Vanderbilt-Ingram Cancer Center SPORE in Breast Cancer (P50CA098131). Flow cytometry experiments were conducted in the Vanderbilt University Medical Center (VUMC) Flow Cytometry Shared Resource which is supported by the Vanderbilt-Ingram Cancer Center (P30 CA68485) and the Vanderbilt Digestive Disease Research Center (DK058404). The VUMC Translational Pathology Shared Resource, supported by a NCI/NIH Cancer Center Support Grant (P30CA068485), performed tissue processing and sectioning. Next Generation Sequencing and digital spatial profiling were completed through the Vanderbilt Technologies for Advanced Genomics (VANTAGE) Core.

## References

1. Zhang M, Deng H, Hu R, Chen F, Dong S, Zhang S, Guo W, Yang W, Chen W. Patterns and prognostic implications of distant metastasis in breast cancer based on SEER population data. Sci Rep. 15(1):26717, 2025.

2. Giaquinto AN, Sung H, Newman LA, Freedman RA, Smith RA, Star J, Jemal A, Siegel RL. Breast cancer statistics 2024. CA Cancer J Clin. 74(6):477–495, 2024.

3. Bergers G, Fendt SM. The metabolism of cancer cells during metastasis. Nat Rev Cancer. 21(3):162–180, 2021.

4. Faubert B, Solmonson A, DeBerardinis RJ. Metabolic reprogramming and cancer progression. Science. 368(6487):eaaw5473, 2020.

5. Karno B, Edwards DN, Chen J. Metabolic control of cancer metastasis: role of amino acids at secondary organ sites. Oncogene. 42(47):3447–3456, 2023.

6. Li Z, Wang L, Ren Y, Huang Y, Liu W, Lv Z, Qian L, Yu Y, Xiong Y. Arginase: shedding light on the mechanisms and opportunities in cardiovascular diseases. Cell Death Discov. 8(1):413, 2022.

7. Niu F, Yu Y, Li Z, Ren Y, Li Z, Ye Q, Liu P, Ji C, Qian L, Xiong Y. Arginase: an emerging and promising therapeutic target for cancer treatment. Biomed Pharmacother. 149:112840, 2022.

8. Hibbs JB, Taintor RR, Vavrin Z. Macrophage cytotoxicity: role for L-arginine deiminase and imino nitrogen oxidation to nitrite. Science. 235(4787):473–476, 1987.

9. Nussler AK, Billiar TR. Inflammation, immunoregulation, and inducible nitric oxide synthase. J Leukoc Biol. 54(2):171–178, 1993.

10. Caldwell RW, Rodriguez PC, Toque HA, Narayanan SP, Caldwell RB. Arginase: a multifaceted enzyme important in health and disease. Physiol Rev. 98(2):641–665, 2018.

11. Mussai F, Egan S, Hunter S, Webber H, Fisher J, Wheat R, McConville C, Sbirkov Y, Wheeler K, Bendle G, Petrie K, Anderson J, Chesler L, De Santo C. Neuroblastoma arginase activity creates an immunosuppressive microenvironment that impairs autologous and engineered immunity. Cancer Res. 75(15):3043–3053, 2015.

12. Fultang L, Gamble LD, Gneo L, Berry AM, Egan SA, De Bie F, Yogev O, Eden GL, Booth S, Brownhill S, Vardon A, McConville CM, Cheng PN, Norris MD, Etchevers HC, Murray J, Ziegler DS, Chesler L, Schmidt R, Burchill SA, Haber M, De Santo C, Mussai F. Macrophage-derived IL1β and TNFα regulate arginine metabolism in neuroblastoma. Cancer Res. 79(3):611–624, 2019.

13. Sousa MSA, Latini FRM, Monteiro HP, Cerutti JM. Arginase 2 and nitric oxide synthase: pathways associated with the pathogenesis of thyroid tumors. Free Radic Biol Med. 49(6):997–1007, 2010.

14. Yang Y, Brenna A, Potenza DM, Sundaramoorthy S, Cheng X, Ming XF, Yang Z. Arginase-II promotes melanoma and lung cancer cell growth by regulating Sirt3-mtROS axis. Front Cell Dev Biol. 13:1528972, 2025.

15. Zhang Y, Qu N, Mei J, Mo C, Wei L, Shen H, Zhou S, Hu J, Luo W, Mo X. Arginase 2 promotes colorectal cancer metastasis via PI3K/AKT pathway activation and regulates tumor immune infiltration. Cancer Med. 15(2):e71567, 2026.

16. Mumenthaler SM, Rozengurt N, Livesay JC, Sabaghian A, Cederbaum SD, Grody WW. Disruption of arginase II alters prostate tumor formation in TRAMP mice. Prostate. 68(14):1561–1569, 2008.

17. Ochocki JD, Khare S, Hess M, Ackerman D, Qiu B, Daisak JI, Worth AJ, Lin N, Lee P, Xie H, Li B, Wubbenhorst B, Maguire TG, Nathanson KL, Alwine JC, Blair IA, Nissim I, Keith B, Simon CM. Arginase 2 suppresses renal carcinoma progression via biosynthetic cofactor pyridoxal phosphate depletion and increased polyamine toxicity. Cell Metab. 27(6):1263–1280.e6, 2018.

18. Ridnour LA, Cheng RYS, Heinz WF, Pore M, Gonzalez AL, Femino EL, Moffat RL, Wink AL, Imtiaz F, Coutinho LL, Butcher D, Edmondson EF, Rangel MC, Wong ST, Lipkowitz S, Glynn SA, Vitek MP, McVicar DW, Li X, Anderson SK, Paolocci N, Hewitt SM, Ambs S, Billiar TR, Chang JC, Lockett SJ, Wink DA. Elevated tumor NOS2/COX2 promotes immunosuppressive phenotypes associated with poor survival in ER– breast cancer. JCI Insight. 10(16):e193091, 2025.

19. Basudhar D, Glynn SA, Greer M, Somasundaram V, No JH, Scheiblin DA, Garrido P, Heinz WF, Ryan AE, Weiss JM, Cheng RYS, Ridnour LA, Lockett SJ, McVicar DW, Ambs S, Wink DA. Coexpression of NOS2 and COX2 accelerates tumor growth and reduces survival in estrogen receptor-negative breast cancer. Proc Natl Acad Sci U S A. 114(49):13030–13035, 2017.

20. Sun L, Wu J, Du F, Chen X, Chen ZJ. Cyclic GMP-AMP synthase is a cytosolic DNA sensor that activates the type I interferon pathway. Science. 339(6121):786–791, 2013.

21. Wu J, Sun L, Chen X, Du F, Shi H, Chen C, Chen ZJ. Cyclic GMP-AMP is an endogenous second messenger in innate immune signaling by cytosolic DNA. Science. 339(6121):826–830, 2013.

22. Burdette DL, Monroe KM, Sotelo-Troha K, Iwig JS, Eckert B, Hyodo M, Hayakawa Y, Vance RE. STING is a direct innate immune sensor of cyclic di-GMP. Nature. 478(7370):515–518, 2011.

23. Ishikawa H, Barber GN. STING is an endoplasmic reticulum adaptor that facilitates innate immune signalling. Nature. 455(7213):674–678, 2008.

24. Li CQ, Trudel LJ, Wogan GN. Nitric oxide-induced genotoxicity, mitochondrial damage, and apoptosis in human lymphoblastoid cells expressing wild-type and mutant p53. Proc Natl Acad Sci U S A. 99(16):10364–10369, 2002.

25. Jaiswal M, Larusso NF, Shapiro RA, Billiar TR, Gores GJ. Nitric oxide-mediated inhibition of DNA repair potentiates oxidative DNA damage in cholangiocytes. Gastroenterology. 120(1):190–199, 2001.

26. Jaiswal M, LaRusso NF, Gores GJ. Nitric oxide in gastrointestinal epithelial cell carcinogenesis: linking inflammation to oncogenesis. Am J Physiol Gastrointest Liver Physiol. 281(3):G626–G634, 2001.

27. Hirota K, Yoshitomi H, Hashimoto M, Maeda S, Teradaira S, Sugimoto N, Yamaguchi T, Nomura T, Ito H, Nakamura T, Sakaguchi N, Sakaguchi S. Preferential recruitment of CCR6-expressing Th17 cells to inflamed joints via CCL20 in rheumatoid arthritis and its animal model. J Exp Med. 204(12):2803–2812, 2007.

28. Bettelli E, Carrier Y, Gao W, Korn T, Strom TB, Oukka M, Weiner HL, Kuchroo VK. Reciprocal developmental pathways for the generation of pathogenic effector TH17 and regulatory T cells. Nature. 441(7090):235–238, 2006.

29. Veldhoen M, Hocking RJ, Atkins CJ, Locksley RM, Stockinger B. TGFβ in the context of an inflammatory cytokine milieu supports de novo differentiation of IL-17-producing T cells. Immunity. 24(2):179–189, 2006.

30. Pang D, Bertocchi A, Powrie F, Pohin M. Contextualizing TH17 cells in cancer. Nat Rev Immunol. 26(5):367–381, 2026.

31. Salazar Y, Zheng X, Brunn D, Raifer H, Picard F, Zhang Y, Winter H, Guenther S, Weigert A, Weigmann B, Dumoutier L, Renauld J, Waisman A, Schmall A, Tufman A, Fink L, Brune B, Bopp T, Grimminger F, Seeger W, Pullamsetti SS, Huber M, Savai R. Microenvironmental Th9 and Th17 lymphocytes induce metastatic spreading in lung cancer. J Clin Invest. 130(7):3560–3575, 2020.

32. Shibabaw T, Teferi B, Ayelign B. The role of Th-17 cells and IL-17 in the metastatic spread of breast cancer: as a means of prognosis and therapeutic target. Front Immunol. 14:1094823, 2023.

33. Nasr Z, Robert F, Porco JA, Muller WJ, Pelletier J. eIF4F suppression in breast cancer affects maintenance and progression. Oncogene. 32(7):861–871, 2013.

34. Singh R, Avliyakulov NK, Braga M, Haykinson MJ, Martinez L, Singh V, Parveen M, Chaudhuri G, Pervin S. Proteomic identification of mitochondrial targets of arginase in human breast cancer. PLoS One. 8(12):e79242, 2013.

35. Johnstone CN, Smith YE, Cao Y, Burrows AD, Cross RSN, Ling X, Redvers RP, Doherty JP, Eckhardt BL, Natoli AL, Restall CM, Lucas E, Pearson HB, Deb S, Britt KL, Rizzitelli A, Li J, Harmey JH, Pouliot N, Anderson RL. Functional and molecular characterisation of EO771.LMB tumours, a new C57BL/6-mouse-derived model of spontaneously metastatic mammary cancer. Dis Model Mech. 8(3):237–251, 2015.

36. Xiong Y, Yepuri G, Forbiteh M, Yu Y, Montani JP, Yang Z, Ming X. ARG2 impairs endothelial autophagy through regulation of MTOR and PRKAA/AMPK signaling in advanced atherosclerosis. Autophagy. 10(12):2223–2238, 2014.

37. Rashid OM, Nagahashi M, Ramachandran S, Dumur CI, Schaum JC, Yamada A, Aoyagi T, Milstien S, Spiegel S, Takabe K. Is tail vein injection a relevant breast cancer lung metastasis model? J Thorac Dis. 5(4):385–392, 2013.

38. Farlik M, Reutterer B, Schindler C, Greten F, Vogl C, Muller M, Decker T. Nonconventional initiation complex assembly by STAT and NF-kappaB transcription factors regulate nitric oxide synthase expression. Immunity. 33(1):25–34, 2010.

39. Geismann C, Grohmann F, Dreher A, Hasler R, Rosenstiel P, Legler K, Hauser C, Egberts JH, Sipos B, Schreiber S, Linkermann A, Hassan Z, Schneider G, Schafer H, Arlt A. Role of CCL20 mediated immune cell recruitment in NF-κB mediated TRAIL resistance of pancreatic cancer. Biochim Biophys Acta Mol Cell Res. 1864(5):782–796, 2017.

40. Ngo KA, Kishimoto K, Davis-Turak J, Pimplaskar A, Cheng Z, Spreafico R, Chen EY, Tam A, Ghosh G, Mitchell S, Hoffman A. Dissecting the regulatory strategies of NF-κB RelA target genes in the inflammatory response reveals differential transactivation logics. Cell Rep. 30(8):2758–2775, 2020.

41. Casero RA, Murray Stewart T, Pegg AE. Polyamine metabolism and cancer: treatments, challenges and opportunities. Nat Rev Cancer. 18(11):681–695, 2018.

42. Simon PS, Sharman SK, Lu C, Yang D, Paschall A V, Tulachan SS, Liu K. The NF-κB p65 and p50 homodimer cooperate with IRF8 to activate iNOS transcription. BMC Cancer. 15:770, 2015.

43. Wright JF, Bennett F, Li B, Brooks J, Luxenberg DP, Whitters MJ, Tomkinson KN, Fitz LJ, Wolfman NM, Collins M, Dunussi-Joannopoulos K, Chatterjee-Kishore M, Carreno BM. The human IL-17F/IL-17A heterodimeric cytokine signals through the IL-17RA/IL-17RC receptor complex. J Immunol. 181(4):2799–2805, 2008.

44. Wu SZ, Al-Eryani G, Roden DL, Junankar S, Harvey K, Andersson A, Thennavan A, Wang C, Torpy JR, Bartonicek N, Wang T, Larsson L, Kaczorowski D, Weisenfeld NI, Uytingco CR, Chew JG, Bent ZW, Chan C, Gnanasambandapillai V, Dutertre C, Gluch L, Hui MN, Beith J, Parker A, Robbins E, Segara D, Cooper C, Mak C, Chan B, Warrier S, Ginhoux F, Millar E, Powell JE, Williams SR, Liu XS, O’Toole S, Lim E, Lundegerg J, Perou CM, Swarbrick A. A single-cell and spatially resolved atlas of human breast cancers. Nat Genet. 53(9):1334–1347, 2021.

45. Arner EN, Rathmell JC. Metabolic programming and immune suppression in the tumor microenvironment. Cancer Cell. 41(3):421–433, 2023.

46. Kim TH, Lim SH, Lee H, Chae YC, Min DS. Metabolic crosstalk among cancer-associated fibroblasts, adipocytes and immune cells as an immunosuppressive tumor microenvironment driver. Exp Mol Med. 58(2):366–381, 2026.

47. Bundgen G, Ulges A, Pietruschka J, Truong-Andrievici N, Klein M, Romaniuk K, Schmitt F, Hagen M, Seebass JG, Zezlina L, Stein L, Probst HC, Distler U, Tenzer S, Lohoff M, Romero-Olmedo AJ, Mei H, Bohn T, Delacher M, Schmidlin T, Gaida MM, Dikic I, Imbusch C, Schild H, Bopp T. Polyamines regulate adaptive antitumor immunity by functional specialization of regulatory T cells. Immunity. 58(8):2019–2034.e11, 2025.

48. Hibino S, Eto S, Hangai S, Endo K, Ashitani S, Sugaya M, Osawa T, Soga T, Taniguchi T, Yanai H. Tumor cell-derived spermidine is an oncometabolite that suppresses TCR clustering for intratumoral CD8+ T cell activation. Proc Natl Acad Sci U S A. 120(24):e2305245120, 2023.

49. Al-Habsi M, Chamoto K, Matsumoto K, Nomura N, Zhang B, Sugiura Y, Sonomura K, Maharani A, Nakajima Y, Wu Y, Nomura Y, Menzies R, Tajima M, Kitaoka K, Haku Y, Delghandi S, Yurimoto K, Matsuda F, Iwata S, Ogura T, Fagarasan S, Honjo T. Spermidine activates mitochondrial trifunctional protein and improves antitumor immunity in mice. Science. 378(6618):eabj3510, 2022.

50. Briukhovetska D, Suarez-Gosalvez J, Voigt C, Markota A, Giannou AD, Schübel M, Jobst J, Zhang T, Dorr J, Markl F, Majed L, Muller PJ, May P, Gottschlich A, Tokarew N, Lucke J, Oner A, Schwerdtfeger M, Andreu-Sanz D, Grunmeier R, Seifert M, Michaelides S, Hristov M, Konig LM, Cadilha BL, Mikhaylov O, Anders HJ, Rothenfusser S, Flavell RA, Cerezo-Wallis D, Tejedo C, Soengas MS, Bald T, Huber S, Endres S, Kobold S. T cell-derived interleukin-22 drives the expression of CD155 by cancer cells to suppress NK cell function and promote metastasis. Immunity. 56(1):143–161.e11, 2023.

51. Edwards DN, Wang S, Kane K, Song W, Kim LC, Ngwa VM, Hwang Y, Ess K, Boothby MR, Chen J. Increased fatty acid delivery by tumor endothelium promotes metastatic outgrowth. JCI Insight. 10(9):e187531, 2025.

52. Uhlen M, Fagerberg L, Hallstrom BM, Lindskog C, Oksvold P, Mardinoglu A, Sivertsson A, Kampf C, Sjostedt E, Asplund A, Olsson I, Edlund K, Lundberg E, Navani S, Szigyarto CA, Odeberg J, Djureninovic D, Takanen JO, Hober S, Alm T, Edqvist P, Berling H, Tegel H, Mulder J, Rockberg J, Nilsson P, Schwenk JM, Hamsten M, von Feilitzen K, Forsberg M, Persson L, Johansson F, Zwahlen M, von Heijne G, Nielsen J, Ponten F. Tissue-based map of the human proteome. Science. 347(6220):1260419, 2015.

