## Supplementary Figures and Tables for "Loss of Arginase 2 Promotes Lung Metastasis in immune-competent hosts via Nitric Oxide Synthase 2-Dependent Th17 Response"

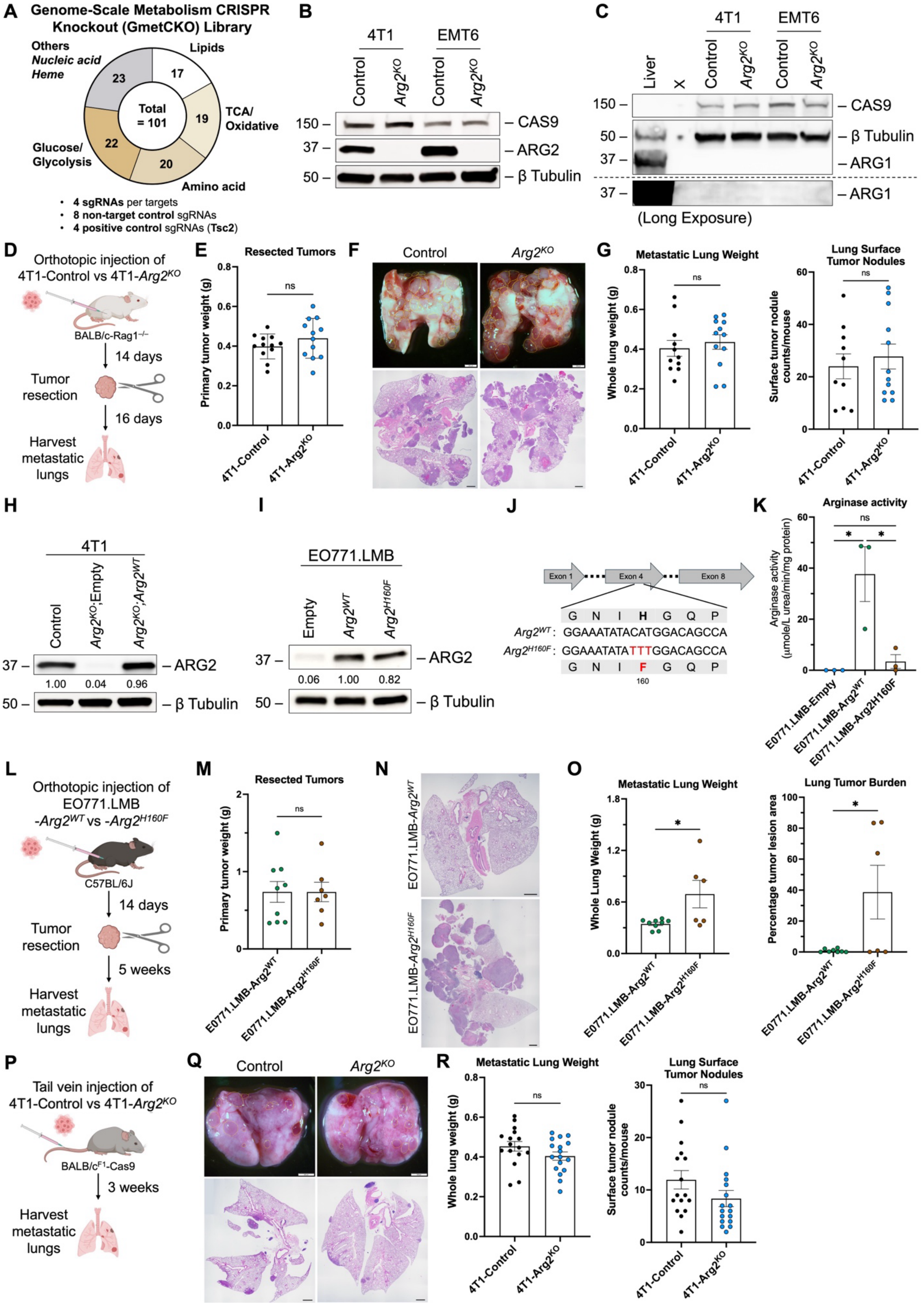
A
Genome-Scale Metabolism CRISPR Knockout (GmetCKO) Library
Others
Nucleic acid
Heme
23
Lipids
17
Total = 101
TCA/ Oxidative
19
Glucose/ Glycolysis
22
20
Amino acid
4 sgRNAs per targets
8 non-target control sgRNAs
4 positive control sgRNAs (Tsc2)
B
4T1
EMT6
Control
Arg2KO
Control
Arg2KO
150 – CAS9
37 – ARG2
50 – β Tubulin
C
4T1
EMT6
Liver
X
Control
Arg2KO
Control
Arg2KO
150 – CAS9
50 – β Tubulin
37 – ARG1
37 – ARG1
(Long Exposure)
D
Orthotopic injection of 4T1-Control vs 4T1-Arg2KO
BALB/c-Rag1–/–
14 days
Tumor resection
16 days
Harvest metastatic lungs
E
Resected Tumors
Primary tumor weight (g)
ns
4T1-Control
4T1-Arg2KO
F
Control
Arg2KO
G
Metastatic Lung Weight
Whole lung weight (g)
ns
4T1-Control
4T1-Arg2KO
Lung Surface Tumor Nodules
Surface tumor nodule counts/mouse
ns
4T1-Control
4T1-Arg2KO
H
4T1
Control
Arg2KO;Empty
Arg2KO;Arg2WT
37 – ARG2
1.00 0.04 0.96
50 – β Tubulin
I
EO771.LMB
Empty
Arg2WT
Arg2H160F
37 – ARG2
0.06 1.00 0.82
50 – β Tubulin
J
Exon 1
Exon 4
Exon 8
G N I H G Q P
Arg2WT : GGAAATATACATGGACAGCCA
Arg2H160F : GGAAATATATTTGGACAGCCA
G N I F G Q P
160
K
Arginase activity
Arginase activity (μmole/L urea/min/mg protein)
ns
*
*
E0771.LMB-Empty
E0771.LMB-Arg2WT
E0771.LMB-Arg2H160F
L
Orthotopic injection of EO771.LMB -Arg2WT vs -Arg2H160F
C57BL/6J
14 days
Tumor resection
5 weeks
Harvest metastatic lungs
M
Resected Tumors
Primary tumor weight (g)
ns
E0771.LMB-Arg2WT
E0771.LMB-Arg2H160F
N
EO771.LMB-Arg2WT
EO771.LMB-Arg2H160F
O
Metastatic Lung Weight
Whole Lung Weight (g)
*
E0771.LMB-Arg2WT
E0771.LMB-Arg2H160F
Lung Tumor Burden
Percentage tumor lesion area
*
E0771.LMB-Arg2WT
E0771.LMB-Arg2H160F
P
Tail vein injection of 4T1-Control vs 4T1-Arg2KO
BALB/cF1-Cas9
3 weeks
Harvest metastatic lungs
Q
Control
Arg2KO
R
Metastatic Lung Weight
Whole lung weight (g)
ns
4T1-Control
4T1-Arg2KO
Lung Surface Tumor Nodules
Surface tumor nodule counts/mouse
ns
4T1-Control
4T1-Arg2KO

**Supplementary Figure 1. Validation of ARG2 expression and function in lung metastasis. A,** The sgRNA library targets metabolic enzymes involved in glucose, nucleotide, heme, amino acid, and lipid metabolism, as well as the tricarboxylic acid (TCA) cycle and oxidative phosphorylation pathways. **B and C,** Western blot analysis of ARG2 (**B**) and ARG1 (**C**) protein expression in 4T1 and EMT6 cells with or without ARG2 deletion. Liver lysate serves as a positive control for ARG1 expression. **D-G,** Validation of ARG2 in regulating spontaneous lung metastasis in *Rag1*$^{-/-}$ mice. **D,** Experimental schematic for the model. **E,** Weights of resected 4T1 primary tumors (n = 12 per group). **F,** Representative gross lung images and H&E-stained lung sections showing metastatic burden. **G,** Whole lung weight and surface tumor nodule counts of harvested lungs. **H and I,** Western blot analysis of ARG2 expression in 4T1 (**H**) and EO771.LMB (**I**) cell lines following Arg2 knockout and re-expression, with quantification of relative intensity by densitometry. **J,** Schematic showing the H160F mutation in ARG2 and the corresponding nucleotide substitution. **K,** Arginase activity was determined in EO771.LMB cells expressing ARG2$^{WT}$ or ARG2$^{H160F}$. **L-O,** *In vivo* spontaneous lung metastasis of EO771.LMB cells expressing ARG2$^{WT}$ or ARG2$^{H160F}$ in C57BL/6J mice. **L,** Experimental schematic of model. **M,** Weights of resected EO771.LMB primary tumors (n = 7 to 9 per group). **N,** Representative H&E-stained lung sections showing metastatic burden. **O,** Whole lung weight and surface tumor nodule counts of harvested lungs. **P-R,** Metastatic lung colonization and outgrowth following tail vein injection of 4T1-Control or 4T1-*Arg2*$^{KO}$ cells. **P,** Experimental schematic of model. **Q,** Representative gross lung images and H&E-stained lung sections showing metastatic burden. **R,** Whole lung weight and surface tumor nodule counts of harvested lungs (n = 16 to 17 per group). Data are presented as mean ± SEM and analyzed using Welch's t-test or one-way ANOVA followed by Tukey's post hoc test. *, $p < 0.05$; **, $p < 0.01$. Scale bar is 200 µm for gross lung image and 1 mm H&E-stained lung sections.

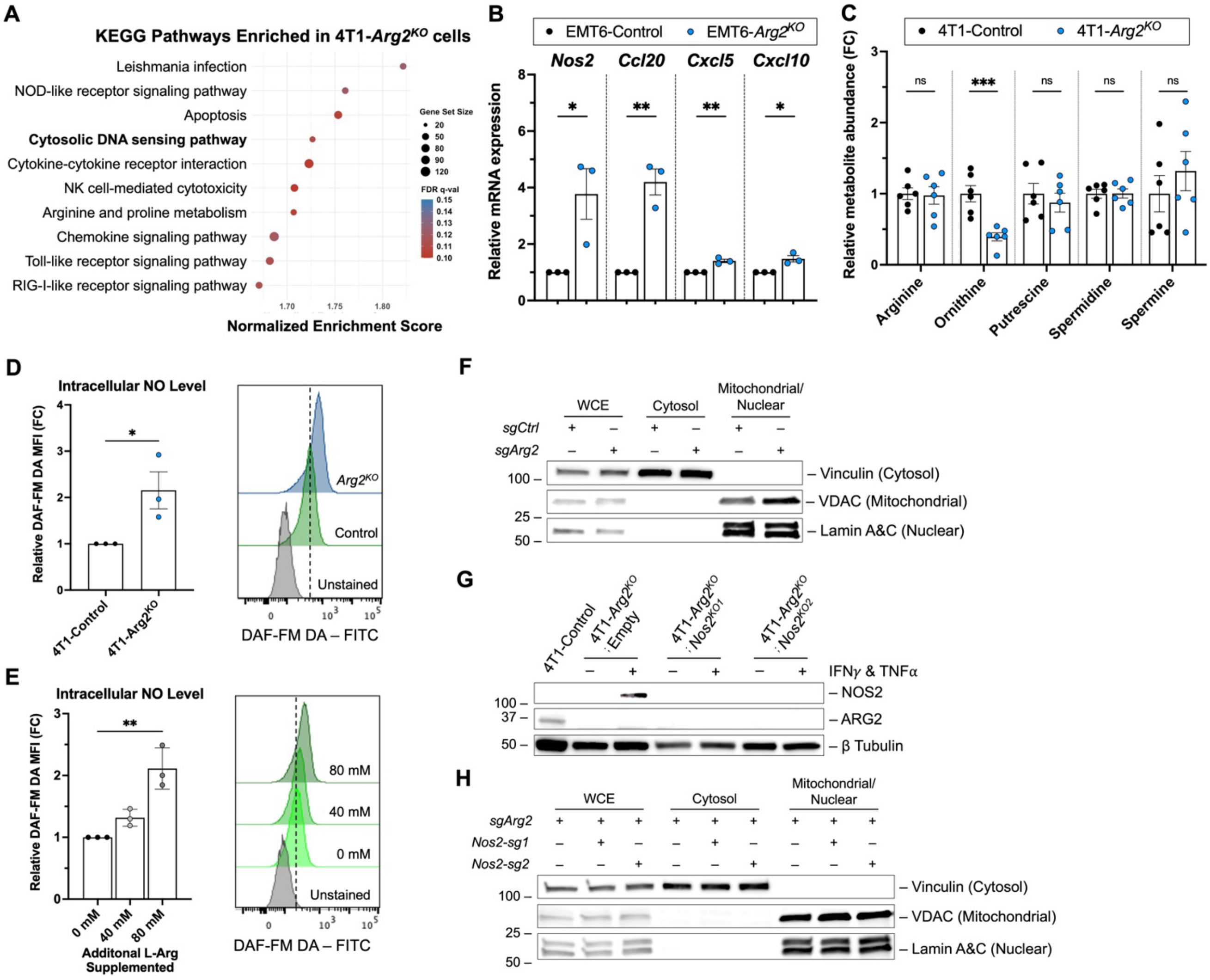


**Supplementary Figure 2. Inflammatory signaling, metabolic alterations, and NOS2-dependent nitric oxide production in ARG2-deficient cells. A,** KEGG pathway enrichment analysis of differentially expressed genes in 4T1-*Arg2^KO^* cells compared with 4T1-Control cells. Gene set size and false discovery rate (FDR) *q*-value are indicated. **B** qRT-PCR analysis of NF-κB target genes (*Nos2, Ccl20, Cxcl5,* and *Cxcl10*) in EMT6-Control and EMT6- *Arg2^KO^* cells (n=3 per group). **C,** Metabolomic analysis of arginine metabolism-related metabolites in 4T1-Control and 4T1-*Arg2^KO^* cells (n=6 per group). **D and E,** Flow cytometry analysis of DAF-FM DA fluorescence to examine intracellular NO level in 4T1-Control and 4T1-*Arg2^KO^* cells (**D**) or 4T1 cells cultured with or without additional L-arginine supplementation (**E**) (n=3 per group). **F,** Western blot analysis of subcellular fractions from control (sgCtrl) or *Arg2^KO^* (sgArg2) 4T1 cells. Vinculin marks the cytosolic fraction, whereas VDAC and lamin A/C mark the mitochondrial and nuclear fractions, respectively. **G,** Western blot analysis confirming deletion of NOS2 in 4T1-*Arg2^KO^;Nos2^KO^* cells. Treatment with TNFα and IFNγ (20 ng/mL each) was used to induce maximal NOS2 expression. **H,** Western blot analysis of subcellular fractions from *Arg2^KO^* (sgArg2) and *Arg2^KO^;Nos2^KO^* 4T1 cells, with fraction markers indicated. Two independent sgNos2 sequences were evaluated. Data are presented as mean ± SEM and analyzed using Welch's t-test *, $p < 0.05$; **, $p < 0.01$; ***, $p < 0.001$.

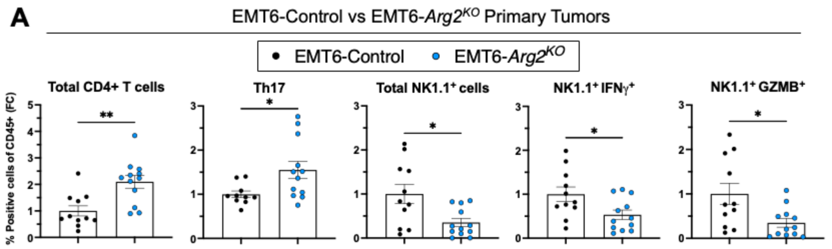


B

| Rank | Gene | log2 fold change | *p* value | Rank Metric Score | Running ES |
|---|---|---|---|---|---|
| 1 | *Cxcl5* | 2.64 | 7.1E-10 | 3.144 | 0.0697 |
| 34 | *Il6* | 2.83 | 1.0E-03 | 1.918 | 0.1099 |
| 43 | *Tgm2* | 0.96 | 4.9E-06 | 1.729 | 0.1477 |
| 125 | *Mmp3* | 0.76 | 4.9E-05 | 1.162 | 0.1673 |
| 147 | *Ccn1* | 0.76 | 1.5E-07 | 1.077 | 0.1896 |
| 370 | *Itgb5* | 0.45 | 1.7E-03 | 0.698 | 0.2668 |
| 473 | *Vegfc* | 0.95 | 5.3E-07 | 0.616 | 0.2726 |
| 514 | *Ccn2* | 0.29 | 3.0E-02 | 0.591 | 0.2827 |
| 574 | *Itga2* | 0.64 | 1.1E-04 | 0.544 | 0.3154 |
| 764 | *Tgfb1* | 0.53 | 1.4E-04 | 1.918 | 0.3561 |
| 1112 | *Serpine2* | 1.06 | 6.0E-06 | 0.353 | 0.3820 |
| 1160 | *Tgfbr3* | 1.08 | 1.2E-05 | 0.339 | 0.3937 |
| 1206 | *Inhba* | 0.49 | 7.3E-04 | 0.327 | 0.3975 |

Table title: **Upregulated EMT-associated genes in 4T1-*Arg2KO* cells (Hallmark – EMT)**

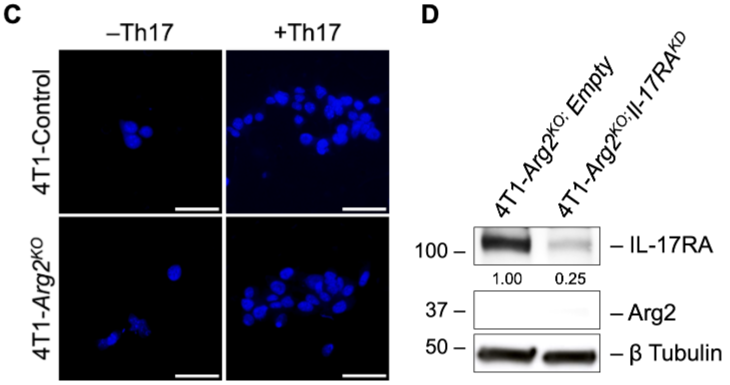


**Supplementary Figure 3. Immune profiling of EMT6 tumors, 4T1 EMT-associated genes, and tumor cell migration in the presence of Th17 cells. A,** Flow cytometric analysis of immune cell populations in EMT6-Control and EMT6-$Arg2^{KO}$ primary tumors. Total $CD4^+$ T cells, $CD4^+$ IL-17$A^+$ (Th17) cells, total NK1.$1^+$ cells, NK1.$1^+$IFN$\gamma^+$ cells, and NK1.$1^+$GZM$B^+$ cells were determined as a percent (%) of $CD45^+$ immune cells. Data are presented as fold change (FC) relative to the control (n=11 to 12 per group). **B,** Ranked list of upregulated EMT-associated genes identified by mRNA sequencing in 4T1-$Arg2^{KO}$ cells from the EMT gene hallmark. $Log_2$ fold change, adjusted $p$ value, rank metric score, and running enrichment score are indicated. **C,** Representative images of transwell migration assay of 4T1-Control or 4T1-$Arg2^{KO}$ cells co-cultured with or without *in vitro*-differentiated Th17 cells. Nuclei of migrated cells were stained with DAPI. Scale bars are 50 µm. **D,** Western blot analysis confirming IL-17RA knockdown in 4T1-$Arg2^{KO}$ cells. Relative densitometry values are shown. Data are presented as mean ± SEM and analyzed using Welch's t-test. *, $p < 0.05$; **, $p < 0.01$.

**Supplementary Table 1. Oligonucleotides, DNA fragments, and plasmids**

**Oligonucleotides**

| **Gene Target** | **Sequence–Forward (5’→3’)** | **Sequence–Reverse (5’→3’)** |
|---|---|---|
| **Quantitative PCR Primers** | | |
| Mouse Arg2 | GGCTGATTGGCAAAAGGCAG | ATCCCCCTACAACAGGGGTT |
| Mouse Rpl37 | CTACCGCAGATTCAGACATGGA | ACCGAACTCTGAACCGATGT |
| Mouse Nos2 | AACTTTACAGGGAGTTGAAGACTGA | CAAATCCAACGTTCTCCGTTCT |
| Mouse Ccl20 | GCAACTACGACTGTTGCCTC | ACATCTTCTTGACTCTTAGGCTGA |
| Mouse Il-6 | TACCACTTCACAAGTCGGAGGC | CTGCAAGTGCATCATCGTTGTTC |
| Mouse Cxcl5 | CCCTACGGTGGAAGTCATAGC | TTAGCTTTCTTTTTGTCACTGCCC |
| Mouse Cxcl10 | GAAATCATCCCTGCGAGCCT | AGTTAAGGAGCCCTTTTAGACCTTT |
| Mouse Mmp3 | ACATGGAGACTTTGTCCCTTTTG | TTGGCTGAGTGGTAGAGTCCC |
| Mouse Mmp13 | GATGACCTGTCTGAGGAAGACC | GCATTTCTCGGAGCCTGTCAAC |
| Mouse mtD-loop1 | AATCTACCATCCTCCGTGAAACC | TCAGTTTAGCTACCCCCAAGTTTAA |
| Mouse mtD-loop2 | CCCTTCCCCATTTGGTCT | TGGTTTCACGGAGGATGG |
| Mouse mt-16S | CACTGCCTGCCCAGTGA | ATACCGCGGCCGTTAAA |
| Mouse mt-Nd4 | AACGGATCCACAGCCGTA | AGTCCTCGGGCCATGATT |
| **CRISPR sgRNA** | | |
| Arg2-sgRNA | caccgCCAAATATTGTGTACATTGG | aaacCCAATGTACACAATATTTGGc |
| LacZ-sgRNA | caccgTGCGAATACGCCCACGCGAT | aaacATCGCGTGGGCGTATTCGCAc |
| Nos2-sgRNA1 | caccgGTGACGGCAAACATGACTTC | aaacGAAGTCATGTTTGCCGTCACc |
| Nos2-sgRNA2 | caccgGAGTTCATCAACCAGTATTA | aaacTAATACTGGTTGATGAACTCc |

**Lentiviral Vectors and Transfection/Transduction Reagents**

| **Vector** | **Source** | **Identifier** |
|---|---|---|
| plentiCRISPRv2-Puro | Addgene | #52961 |
| plentiCRISPRv2-Blast | Addgene | #98293 |
| plenti-CMV-Blast | Addgene | #17486 |
| pCMV-VSV-G | Addgene | #8454 |
| psPAX2 | Addgene | #12260 |
| Lipofectamin™ 3000 Transfection Reagent | Thermo Fisher Scientific | #L3000-015 |
| Polybrene | Sigma-Aldrich | #TR-1003-G |

**gBlock DNA**

| **Gene Target** | **Sequence (5’→3’)** |
|---|---|
| $Arg2^{WT}$ | cctccatagaagacaccgacTCTAGAGCCACCATGTTCCTGAGGAGCAGCGCCTCCCGTCTCC<br>TCCACGGGCAAATTCCTTGCGTCCTGACGAGATCCGTCCACTCTGTAGCTATAGTC<br>GGAGCCCCTTTCTCTCGGGGACAGAAGAAGCTAGGAGTGGAATATGGTCCAGCTG<br>CCATTCGAGAAGCTGGCTTGCTGAAGAGGCTCTCCAGGTTGGGATGCCACCTAAA<br>AGACTTTGGAGACTTGAGTTTTACTAATGTCCCACAAGATGATCCCTACAATAATC<br>TGGTTGTGTATCCTCGTTCAGTGGGCCTTGCCAACCAGGAACTGGCTGAAGTGGTT<br>AGTAGAGCTGTGTCAGGTGGCTACAGCTGTGTCACCATGGGAGGAGACCACAGCC<br>TGGCAATAGGTACCATTATCGGTCACGCCCGGCACCGCCCAGATCTCTGTGTCATC<br>TGGGTTGATGCTCATGCGGACATTAATACACCTCTCACCACTGTATCTGGAAATAT<br>ACATGGACAGCCACTTTCCTTTCTCATCAAAGAACTACAAGACAAGGTACCACAA<br>CTGCCAGGATTTTCCTGGATCAAACCTTGCCTCTCTCCgCCcAATATaGTcTACATTG<br>GCCTGAGAGATGTGGAGCCTCCTGAACATTTTATTTTAAAGAATTATGACATCCA<br>GTATTTTTCCATGAGAGAGATTGATCGACTTGGGATCCAGAAGGTGATGGAACAG<br>ACATTTGATCGGCTGATTGGCAAAAGGCAGAGGCCAATCCACCTGAGTTTTGACA<br>TTGATGCATTTGACCCTAAACTGGCTCCAGCCACAGGAACCCCTGTTGTAGGGGG<br>ATTAACCTACAGAGAAGGAGTGTATATTACTGAAGAAATACATAATACAGGGTTG<br>CTGTCAGCTCTGGATCTTGTTGAAGTCAATCCTCATTTGGCCACTTCTGAGGAAGA<br>GGCCAAGGCAACAGCCAGACTAGCAGTGGATGTGATTGCTTCAAGTTTTGGTCAG<br>ACAAGAGAAGGAGGACACATTGTCTATGACCACCTTCCTACTCCTAGTTCACCAC<br>ACGAATCAGAAAATGAAGAATGTGTGAGAATTTAGgtcgacaatcaacctctggattacaa |

| | |
|---|---|
| $Arg2^{H160F}$ | cctccatagaagacaccgacTCTAGAGCCACCATGTTCCTGAGGAGCAGCGCCTCCCGTCTCC<br>TCCACGGGCAAATTCCTTGCGTCCTGACGAGATCCGTCCACTCTGTAGCTATAGTC<br>GGAGCCCCTTTCTCTCGGGGACAGAAGAAGCTAGGAGTGGAATATGGTCCAGCTG<br>CCATTCGAGAAGCTGGCTTGCTGAAGAGGCTCTCCAGGTTGGGATGCCACCTAAA<br>AGACTTTGGAGACTTGAGTTTTACTAATGTCCCACAAGATGATCCCTACAATAATC<br>TGGTTGTGTATCCTCGTTCAGTGGGCCTTGCCAACCAGGAACTGGCTGAAGTGGTT<br>AGTAGAGCTGTGTCAGGTGGCTACAGCTGTGTCACCATGGGAGGAGACCACAGCC<br>TGGCAATAGGTACCATTATCGGTCACGCCCGGCACCGCCCAGATCTCTGTGTCATC<br>TGGGTTGATGCTCATGCGGACATTAATACACCTCTCACCACTGTATCTGGAAATAT<br>AtttGGACAGCCACTTTCCTTTCTCATCAAAGAACTACAAGACAAGGTACCACAACT<br>GCCAGGATTTTCCTGGATCAAACCTTGCCTCTCTCCgCCcAATATaGTcTACATTGG<br>CCTGAGAGATGTGGAGCCTCCTGAACATTTTATTTTAAAGAATTATGACATCCAGT<br>ATTTTTCCATGAGAGAGATTGATCGACTTGGGATCCAGAAGGTGATGGAACAGAC<br>ATTTGATCGGCTGATTGGCAAAAGGCAGAGGCCAATCCACCTGAGTTTTGACATT<br>GATGCATTTGACCCTAAACTGGCTCCAGCCACAGGAACCCCTGTTGTAGGGGGAT<br>TAACCTACAGAGAAGGAGTGTATATTACTGAAGAAATACATAATACAGGGTTGCT<br>GTCAGCTCTGGATCTTGTTGAAGTCAATCCTCATTTGGCCACTTCTGAGGAAGAGG<br>CCAAGGCAACAGCCAGACTAGCAGTGGATGTGATTGCTTCAAGTTTTGGTCAGAC<br>AAGAGAAGGAGGACACATTGTCTATGACCACCTTCCTACTCCTAGTTCACCACAC<br>GAATCAGAAAATGAAGAATGTGTGAGAATTTAGgtcgacaatcaacctctggattacaa |

| **Supplementary Table 2. Western blot and flow cytometry antibodies** | | |
|---|---|---|
| **Reagent or Resource** | **Source** | **Identifier** |
| **Western Blot Antibodies** | | |
| ARG2 | Abcam | #ab228963 |
| ARG1 | Cell Signaling Technology | #93668 |
| β-tubulin | Sigma-Aldrich | #T4026 |
| Cas9 | Cell Signaling Technology | #14697 |
| NOS2 (iNOS) | Cell Signaling Technology | #13120 |
| NF-κB-p65 | Cell Signaling Technology | #8242 |
| Phospho-NF-κB-p65 (S536) | Cell Signaling Technology | #3033 |
| STING | Cell Signaling Technology | #50494 |
| Phospho-STING (S365) | Cell Signaling Technology | #9234 |
| Vinculin | Cell Signaling Technology | #13901 |
| VDAC | Cell Signaling Technology | #4661 |
| Lamin A+C | Abcam | #ab169532 |
| Anti-rabbit IgG HRP-linked | Cell Signaling Technology | #7074 |
| Anti-mouse IgG HRP-linked | Cell Signaling Technology | #7076 |
| **Flow Cytometry Antibodies and Viability Dye** | | |
| CD45–APC-Cy7 (30-F11) | BD Biosciences | #557659 |
| TCRβ–PE-Cy7 (H57-597) | Cytek Biosciences | #60-5961 |
| CD8–redFluor 710 (53-6.7) | Cytek Biosciences | #80-0081 |
| CD4–PE/Dazzle 594 (GK1.5) | BioLegend | #100455 |
| NK1.1–Brilliant Violet 605 (PK136) | BioLegend | #108739 |
| IL-17A–PerCP/Cyanine5.5 (TC11-18H10.1) | BioLegend | #506920 |
| IFNγ–violetFluor 450 (XMG1.2) | Cytek Biosciences | #75-7311 |
| Granzyme B–PE (NGZB) | Thermo Fisher Scientific | #12-8898-82 |
| TNFα–APC (MP6-XT22) | Thermo Fisher Scientific | #17-7321-82 |
| Ghost Dye Violet 510 | Cytek Biosciences | #13-0870 |
| Anti-CD16/32 Mouse Fc Block | Cytek Biosciences | #70-0161 |
